# Gonadulins, the fourth type of Insulin-related peptides in Decapods

**DOI:** 10.1101/2020.03.19.998484

**Authors:** Jan A. Veenstra

**Author notes:** Corresponding Author: Jan A. Veenstra INCIA UMR 5287 CNRS, Université de Bordeaux, allée Geoffroy St Hillaire, CS 50023, 33 615 Pessac Cedex, France.

## Abstract

Insulin and related peptides play important roles in the regulation of growth and reproduction. Until recently three different types of insulin-related peptides had been identified from decapod crustaceans. The identification of two novel insulin-related peptides from *Sagmariasus verreauxi* and *Cherax quadricarinatus* suggested that there might a fourth type. Publicly available short read archives show that orthologs of these peptides are commonly present in these animals. Most decapods have two genes coding such peptides, but *Penaeus* species have likely only one and some palaemonids have three. Interestingly, expression levels can vary more than thousand-fold in the gonads of *Portunus trituberculatus*, where gonadulin 1 is expressed by the testis and gonadulin 2 by the ovary. Although these peptides are also expressed in other tissues, the occasionally very high expression in the gonads led to them being called gonadulins.

## Introduction

Neuropeptides and neurohormones originated early during evolution and are well conserved between decapods and insects (*e.g.* Veenstra, 2016). In several cases their functions are also the same or at least similar, this is *e.g.* the case of crustacean cardioactive peptide (CCAP) that plays crucial roles during molting in both decapods and insects (Webster et al., 2013). Other peptides on the other hand likely have very different functions. EFLamide for example is very abundantly expressed and effects many different tissues in decapods (Dickinson et al., 2019), but is present in only some insect species and when present expressed in very few neurons (Veenstra and Šimo, 2020; Kotwica-Rolinska et al., 2020). Allatotropin on the other hand is commonly present in insects but absent from decapods (Veenstra, 2016).

Insulin and related peptides seem to be present in all animals and have been extensively studied in insects where they are important growth hormones. They have been particularly well investigated in *Drosophila* (see *e.g*. Brogiolo et al., 2001; Nässel and Vanden Broeck, 2016), but also in other species such as the silk worm *Bombyx mori* (Mizoguchi and Okamoto, 2013). The best known crustacean insulin-related peptide is the one secreted by the androgenic gland. This gland has been found in all Malacostraca and secretes a hormone that is able to change the sex of animals from female to male as first demonstrated in a amphipod by transplanting those glands (Charniaux-Cotton, 1954). After the initial identification of the hormone made by these glands from an isopod as an insulin-related peptide (Martin et al., 1999) it has also been identified from several decapods (*e.g.* Ventura et al., 2009, 2011; Chung et al., 2011). However the physiological relevance of the other insulin-related peptides in crustaceans remains unclear.

Both growth and sex determination are essential for the survival of a species and hence one would expect that the way these processes are regulated is under significant evolutionary pressure and might be conserved between insects and crustaceans. The insect insulin-like growth hormones and the decapod Insulin-like Androgenic Gland hormone, that will be called androgenin from here on, are evolutionarirly related and I previously suggested that perhaps the androgenins might be orthologs of *Drosophila* insulin-related peptide 8 (dilp 8), one of the *Drosophila* insulin-related peptides (Veenstra, 2016). Unlike most insulin-related peptides that stimulate a receptor tyrosine kinase (RTK) dilp 8 acts through a leucine-rich repeat G-protein coupled receptor (LGR; Vallejo et al., 2015; Gontijo and Garelli, 2018). Dilp8 is made by the imaginal disks, that are only present in holometabolous insects, as well as the ovary and testis. The specific LGR (*Drosophila* LGR3) that is the dilp 8 receptor has orthologs in many other arthropods, including decapods (Veenstra, 2016), meaning that dilp 8 orthologs must be common. Nevertheless, the primary amino acid sequences of dilp 8 and the decapod androgenins are very different. The sequence of dilp 8 is also very poorly conserved within insects. When only three different types of insulin-like peptides had been identified in decapods it seemed an attractive hypothesis that dilp 8 might be an ortholog of androgenin (Veenstra, 2016). After all, the other two are clear orthologs of dilp 7, a relaxin-like peptide, and the insect insulin-related peptides that stimulate growth (Brogiolo et al., 2001; Nässel and Vanden Broeck, 2016). However, soon afterwards two novel decapod insulin-like peptide precursors were identified from *Sagmariasus verreauxi* and *Cherax quadricarinatus* that look more similar to dilp 8 than androgenin (Chandler et al., 2017). This suggests that decapods do not have three but at least four different types of insulin-like peptides and that dilp 8 is probably not an ortholog of androgenin. This provided stimulation to exploit the large amount of genomic and especially trancriptomic sequences available for decapods for additional clues as to the presence and distribution of insulin-related peptides in decapods. I here report that the peptides initially described from *S. verreauxi* and *C. quadricarinatus* belong to a novel type of decapod insulin-like peptides and in many species is produced by two and in some species even three paralogous genes, one or two of which seem to be rarely expressed. However, under particular conditions, that remain to be identified, the transcripts for these peptides are occasionally made in very large numbers by the gonads, at least in the crab *Portunus trituberculatus*. As the gonads seem to be an important site of expression of these peptides, they have been baptized gonadulins.

## Materials and Methods

I analyzed transcriptome and genome SRAs using the tools from the SRAtoolkit (https://trace.ncbi.nlm.nih.gov/Traces/sra/sra.cgi?view=software), Trinity (Grabherr et al., 2011) and Artemis (Rutherford et al., 2000) on the publicly available decapod genomes and draft genomes, as well as a large number of transcriptome (Table S1) and genome SRAs. Details of the methods that were used have been described in more detail elsewhere (Veenstra and Khammassi, 2017; Veenstra, 2019). A list of all SRAs used can be found in the supplementary data. Draft genome assembliess for the following species have been used: *Penaeus vannamei* (Zhang et al., 2019), *P. monodon* (Van Quyen et al., 2020), *Palaemon carinicauda* (Yuan et al., 2017), *Procambarus virginalis* (Gutekunst et al., 2018), *Cherax quadricarinatus* (unpublished, but available at https://www.ncbi.nlm.nih.gov/assembly/GCA_009761615.1/), and two assemblies each for *Eriocheir sinensis* (Song et al., 2016; Tang et al., 2020b) and *Portunus trituberculataus* (https://www.ncbi.nlm.nih.gov/assembly/GCA_008373055.1/; Tang et al., 2020a). Most of these genome assemblies were localized using https://www.ncbi.nlm.nih.gov/genome/?term=decapoda, while the the first *E. sinensis* genome was downloaded from http://gigadb.org/dataset/100186, and the second *P. trituberculatus* genome from http://gigadb.org/dataset/100678.

Decapod genomes tend to be large. A number of draft decapod genomes used Illumina technology for the assembly and these have usually relatively short contigs and no or poor scaffolds. For example, the *Cherax quadricarinatus* genome lacks the second coding exon from the gonadulin 1 gene. The sequence of this exon was obtained by using Trinity on individual reads from genomic SRAs. For the genome assembly of *Penaeus vannamei* and the second assemblies for *Portunus trituberculatus* and *Eriocheir sinensis* PacBio sequencing technology was used. This allows the construction of large scaffolds, but it often suffers from significant numbers of indels, which in some genome stretches are numerous and can make analysis difficult. In some cases certain genes did not make it into the final assembly, *e.g.* the *Portunus* RTK4 gene. Nevertheless these genome assemblies are useful for elucidating the exon-intron structures of genes. Other genomic sequences were recovered from several genomic resources created to establish the phylogenetic relations within the decapods (Wolfe et al., 2019). This concerns data from *Coenobita clypeatus, Stenopus hispidus, Fantapenaeus duorarum, Ocypode quadrata, Menippe nodifrons,* and *Procamabarus clarkii*. These genomic SRAs were used in combination with Trinity to deduce the exon-intron structures of several genes. In some cases, such as that of the *Stenopus hispidus* gonadulin gene, this was done after determining its coding sequence from transcriptome data, while in other species the transcripts from related species were used. For example, the partial *Coenobita clypeatus* genes structures were found in the genomic SRA using the gonadulin precursor deduced from transcriptome SRAs of *Paralithodes camtschaticus* (a species from the same infraorder), while the *Palaemon serratus* and *Macrobrachium* sequences allowed the identification of the *P. carinicauda* genes. Neither of the two *Eriocheir* genome assemblies contain the gonadulin 1 gene, so the exon-intron structure of this gene was also determined from genomic SRAs.

### Phylogenetic trees

The available decapod RTK sequences (spreadsheet1 of supplementary data) were aligned with Clustal omaga (Sievers et al., 2011) and then manually inspected with Seaview (Gouy et al., 2010). The latter program was also used to trim poorly aligned sequences. Fasttree (Price et al., 2010) was employed to make a phylogenetic tree using the ./FastTreeDbl command with the -spr 4 -mlacc 2 -slownni -pseudo options. A second tree was made using exclusively the tyrosine kinase domains (Fig. S2). The phylogenetic tree of the LGRs was made using exclusively the transmembrane regions of these receptors using the same methods. The complete precursors of a number of decapod insulin-related peptides together with the precursors of the eight *Drosophila* insulin-related peptides were aligned with clustal omega and then used with Fasttree to make a tree (Fig. S1). This tree can not be considered a phylogenetic tree as the alignment is far too poor, as illustrated by the low branch probabilities separating the major groups. Nevertheless, it clusters similar peptide sequences and nicely reveals the four major types of decapod insulin-like peptides as well as to which decapod peptides *Drosophila* ilp 8 is most similar in structure. When more than one precursor from the same genus was available, only one was used to eliminate bias as much as possible.

### Sequence logos

As an alternative way to show sequence similarities of the different insulin-related precursors sequence logos were made of those parts of the precursor sequences that show significant conservation between different species using https://weblogo.berkeley.edu/logo.cgi.

### Expression

Expression data were obtained by using the parts of the transcripts that code for the various peptides and receptors (thus without untranslated 5’- and 3’-sequences) as query for the blastn_vdb command from the SRAtoolkit to look for individual reads in the various transcriptome SRAs. Raw counts and results are expressed as reads per million and reported in spreadsheet2 of supplementary data. In the case of the SRAs from *Portunus,* data for a partial vitellogenin receptor sequence was also included in order to show the likely mix up of the ovary and testis SRAs SRR1920180 and SRR1920182.

## Results

This manuscript describes the gonadulins and their putative receptors, however other decapod insulin-like peptides were also identified in an attempt to establish where all the different insulin-related peptides and their putative receptors are expressed. Although gonadulins were identified in many decapod species, identification of their putative receptors and the expression of these genes was limited to eight species. These are *Penaeus vannamei, P. monodon, Macrobrachium nipponense, Cherax quadricarinatus, Procambarus clarkii, Portunus trituberculatus, Carcinus maenas* and *Eriocheir sinensis* (spreadsheet1, spreadsheet2).

Primary amino acid sequences of arthropod insulin-related peptides are notoriously variable, the best conserved sequence features are the the six cysteine residues responsible for the three disulfide bridges. Even that character is not constant, as some peptides have eight cysteine residues and are thus predicted to have four disulfide bridges. This includes some of the decapod gonadulins (Figs 1, 2) as well as some androgenins. It is worth mentioning that, surprisingly, some of the decapod relaxins are predicted to have seven cysteine residues in the propeptide (Table S2). The high primary amino acid sequence variability precludes making reliable phylogenetic trees using these sequences. However, such trees, which are better considered similarity trees, do show that the decapod insulins cluster in four groups and also that *Drosophila* ilp 8 is more similar to the gonadulins than to any of the other decapod insulins (Fig. S1). The primary amino acid residues immediately surrounding the cysteines are often somewhat better conserved, at least in the decapod insulins and relaxins, but hardly or not at all in the androgenis and gonadulins (Fig. 1). It is thus impossible to predict how the different peptides are related based on their primary amino acid sequences, even though the B-chain sequence logos may suggest that androgenin is more similar to relaxin than to the other two peptides (Fig. 1). Deduced amino acid sequences for a number of decapod insulin-related peptides are shown in figures 1 and 2 and more sequences are presented in the supplementary data (Tables S2-S6).

**Fig. 1.**
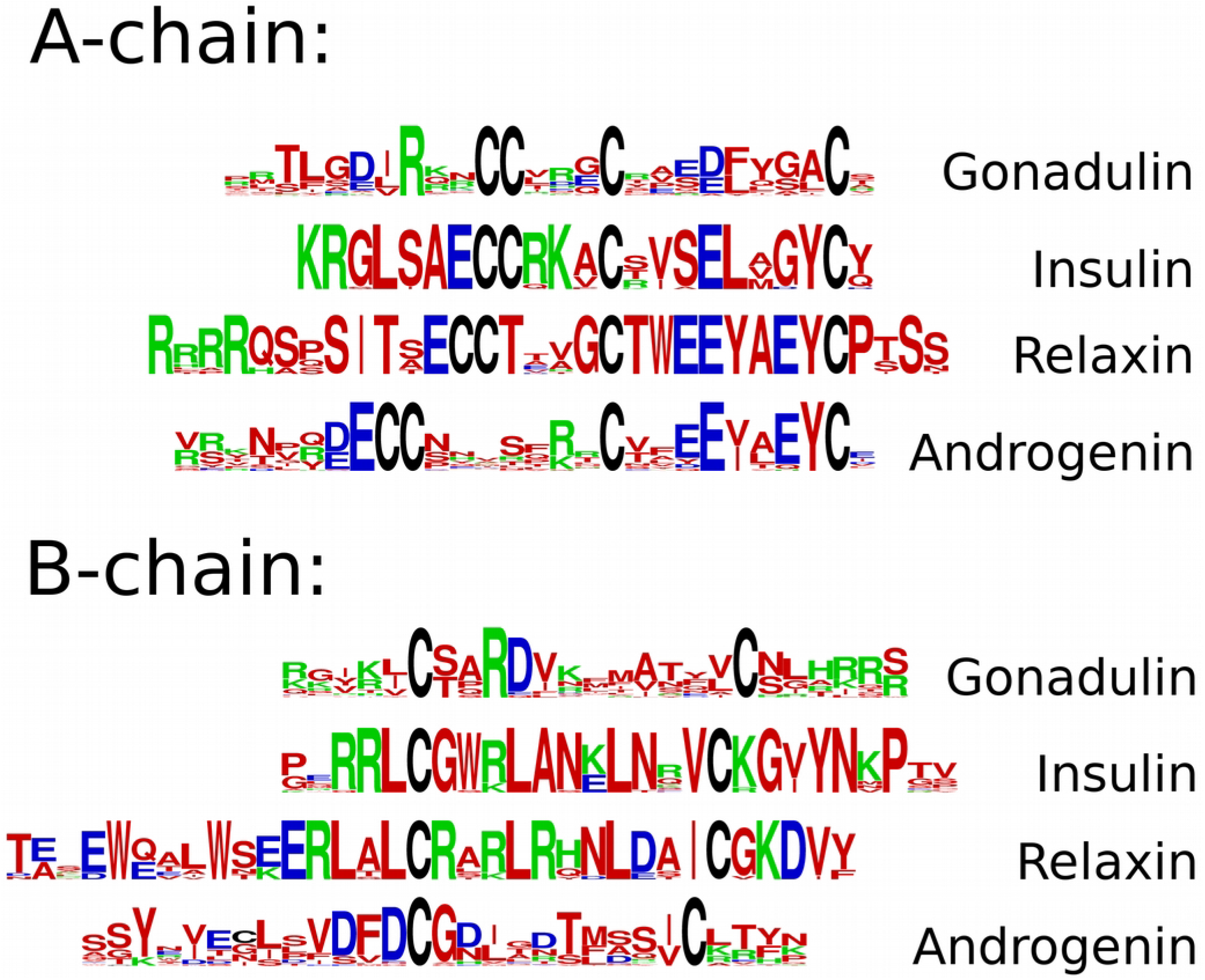
Sequence logos for the A- and B-chains of gonadulin (gon), insulin (ins), relaxin (rel) and androgenin (and) from decapods. Note the conserved arginine located three residues before the first cysteine of the A-chain and the absence of the glutamic acid residue immediately before this cysteine residue that is always present in the other insulin-related peptides. Also note that both insulin and relaxin have the best conserved sequences. Sequences are provided in Tables S2-S6.

**Fig. 2.**
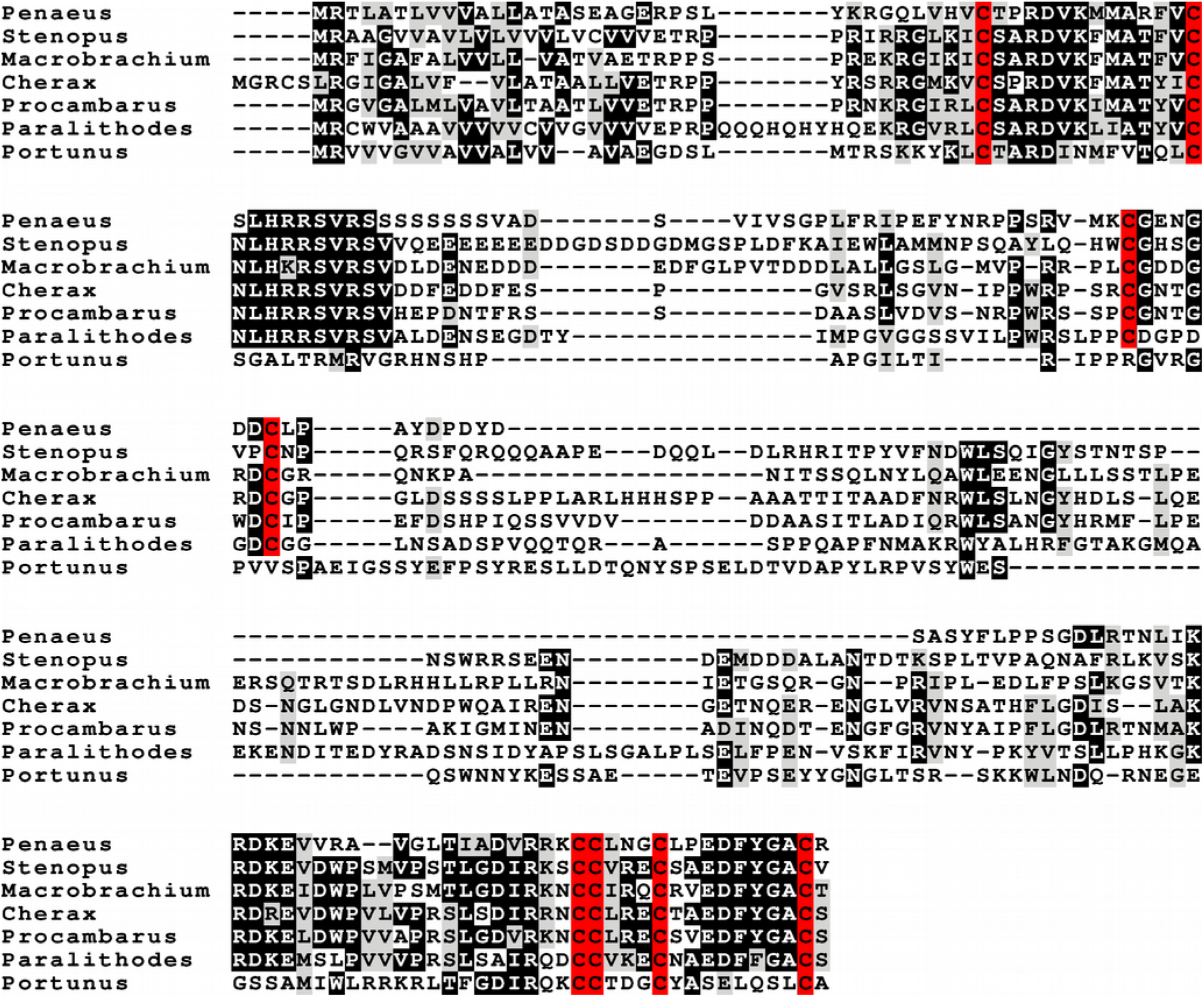
Sequence alignment of a number of gonadulin 1 sequences representing different decapod infraorders: *Penaeus vannemei* (from Trinity), *Stenopus hispidus* (Trinity), *Macrobrachium nipponense* (GHMG01073270.1), *Cherax quadricarinatus* (AIU40993.1), *Procambarus clarkii* (GARH01003670.1), *Paralithodes camtschaticus* (GHJC01030171.1) and *Portunus trituberculatus* (GFFJ01045966.1). Additional gonadulin 1 sequences can be found in table S3. Cysteine resiudes are highlighted in red and show the typical pattern associated with the disulfide bridges in insulin. In most species there is an additional pair of cysteines, but not in *P. trituberculatus.* Sequences are provided in spreadsheet1.

### Gonadulins

Interestingly, many decapods may have two different types of gonadulins and the palaemonids have even three. On the other hand only a single gonadulin was found in three penaeid species or the single species from the Stenopodidae for which data is available. In the first case there is a large amount of data, including genomes for *Penaeus vannamei* and *P. monodon*, as well as a genomic SRA for *Fantapenaeus duorarum* that covers its genome 27-fold. In none of these data sets evidence for a second gonadulin gene could be found, suggesting that the penaeids have only one such gene. For *Stenopus hispidus* there is a genomic SRA with 14-fold coverage, suggesting that it too likely has only a single gonadulin gene.

The different gonadulin genes can be distinguished in three different types, with the last type only found in palaemonidae; they will be referred to as gonadulin 1, gonadulin 2 and gonadulin 3 (Figs 2,3; Tables S3,S4,S5). Of these gonadulin 1 is most widely expressed as detailed below and has also the best conserved primary amino acid sequence. In most species gonadulin 1 has eight cysteine residues rather than the characteristic six typical of insulin (Fig. 2), however gonadulin 2 and 3, which are quite similar to one another, have only six cysteine residues and this is also the case for gonadulin 1 from Brachyura. The sequences corresponding to the putative connecting peptides of gonadulin are often poorly conserved, apart from the piece surrounding the two additional cysteine residues in gonadulin 1. The parts of the gonadulin precursors corresponding to the A-and B-chains are much better conserved, but are still quite variable (Figs 1-3). One structural character that seems to be typical of the gonadulins is the presence of one or more dibasic amino acid residues preceding the first cysteine residue in the A-chain while acidic amino acid residues are absent there. In particular the third residue preceding the first cysteine is always an arginine (Figs 1-3). In contrast, insulin (Table S6), relaxin and androgenin always have an aspartic residue immediately preceding this cysteine (Fig. 1) and usually lack that arginine residue.

**Fig. 3.**
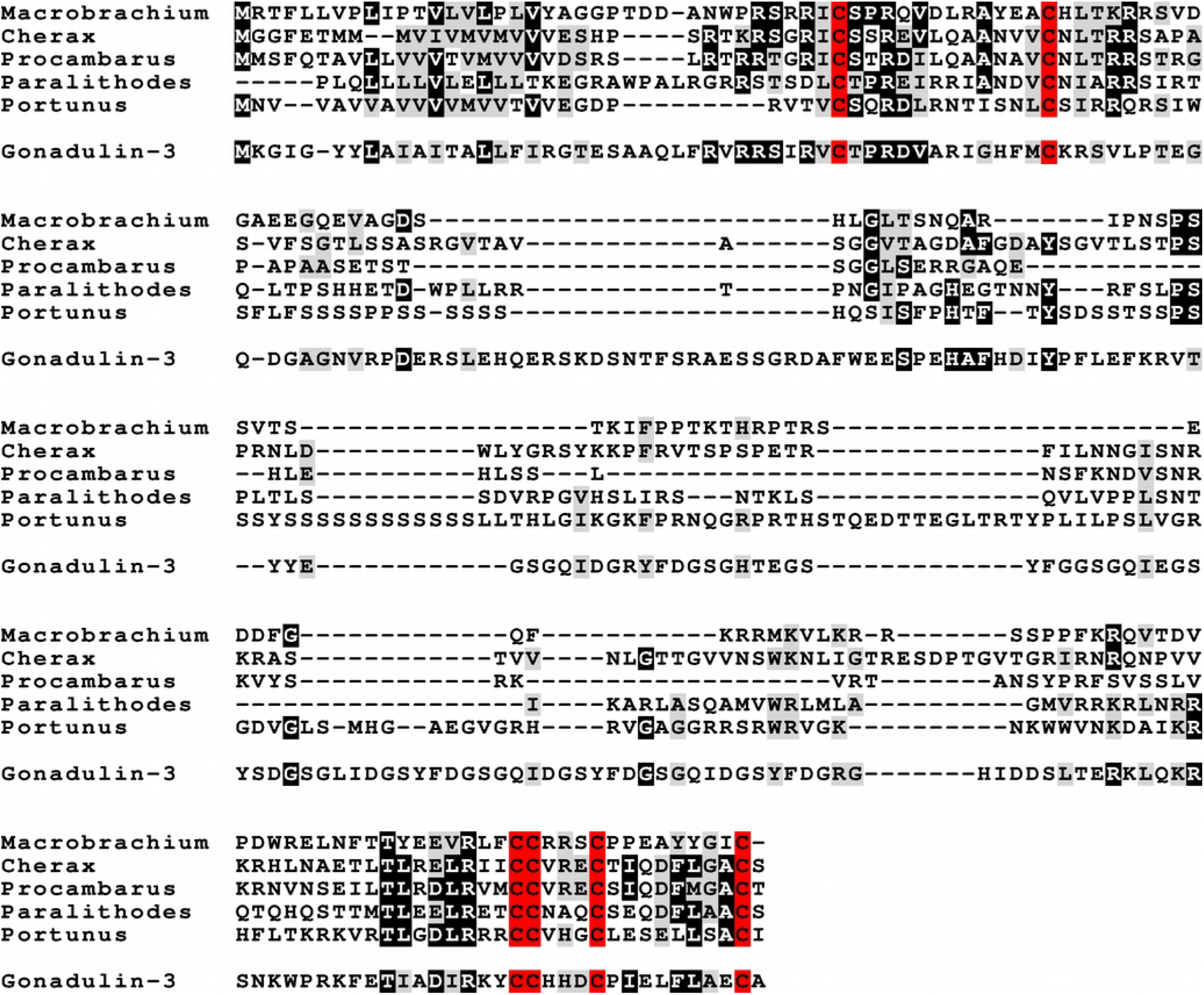
Sequence alignment of a number of gonadulin 2 sequences representing different decapod infraorders together with a gonadulin 3 sequence. Note that gonadulin 2 has the cysteine residues that characterize insulin, but lack the additional disulfide bridge of gonadulin 1. The species are: *Macrobrachium nipponense* (Trinity), *Cherax quadricarinatus* (from genome), *Procambarus clarkii* (from genome SRA), *Paralithodes camtschaticus* (GHJC01008804.1) and *Portunus trituberculatus* (Trinity). These are the same species as in Fig. 1, except for *P. vannamei* and *S. hispidus*, which appear to lack a gonadulin 2 gene. The gonadulin 3 gene is from *M. rosenbergii* and was obtained using Trinity. Sequences are provided in tables S3 and S4.

It is not clear at all how these gonadulin precursors are processed. If they are expressed by the ovary or testes, they are likely to be secreted through the constituitive pathway that has furin but is expected to lack the typical endocrine convertases. On the other hand gonadulin 1 is expressed in the eyestalk, where such enzymes are present. Furin, is likely involved in the processing of these precursors, but its precise cleavage preferences are unknown even in vertebrates, although we do know that they differ between insects and vertebrates (Cano-Monreal et al., 2010). This makes predictions as where convertase(s) may cleave the gonadulin precursors impossible.

### Gene structure

The availability of several (draft) genome assemblies as well as a number of genome SRAs makes it possible to identify the individual exons of the various gonadulin genes. The gonadulin 1 genes for which a gene structure could be deduced have four, five or six coding exons, while the gonadulin 2 genes have consistently two. The single *Penaeus* gene has three coding exons and represents perhaps an ancestral gene that was duplicated. The single *Stenopus* gene has four coding exons and encodes a gonadulin 1. The difference in number of coding exons is not the only difference between the gonadulin 1 and 2 genes. For those species where this can be determined gonadulin 1 genes stretch out over thousands of base pairs and the intron between the two first coding exons accounts for several thousand nucleotides. Although this latter point is most convincingly established in *Portunus*, due to the use of PacBio sequencing, the genome drafts for *Procambarus virginalis* and *Cherax* also show significant sizes for this intron. The gonadulin 2 gene on the other hand has a small intron between its two coding exons; so small in fact, that Trinity sometimes produces a single contig from genomic DNAseq reads containing the two coding exons. The third gonadulin gene, which has only been detected in palaemonids, has two coding exons, like the second gene and the predicted peptides are also similar. The gonadulin precursor sequences of the Palaemonid species are sufficiently similar to allow reconstruction of the various coding exons of the three *P. carinicauda* genes (Fig. 4).

**Fig. 4.**
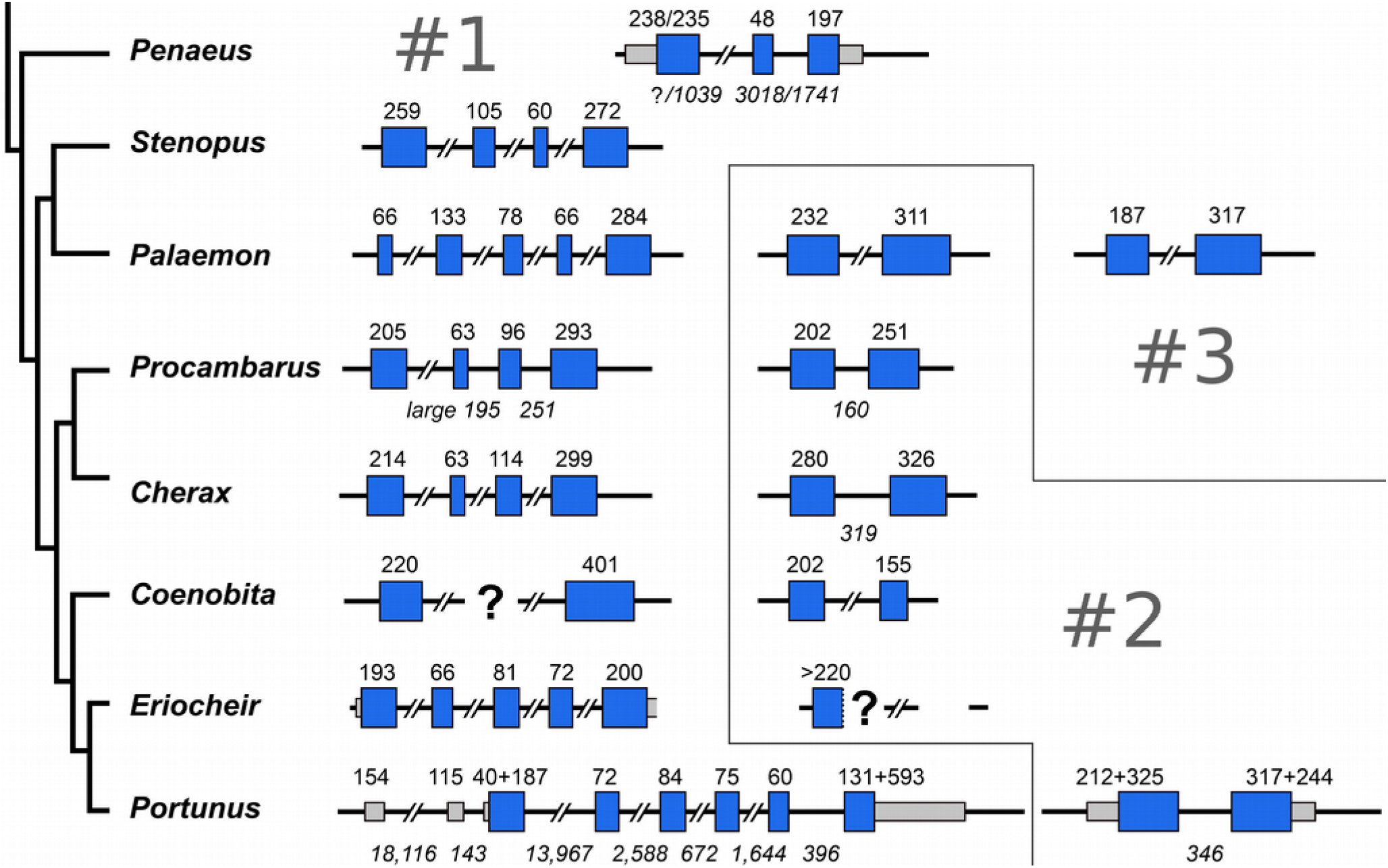
*Configuration of the various decapod gonadulin genes. The peptides coded by the genes from Penaeus vannamei* and *P. monodon* are most similar to gonadulin 1, as is also evident from the primary sequences of the peptides. Note that the gonadulin 2 genes have much smaller introns than the ones that code gonadulin 1. The dendrogram on the left side of the figure reproduce the most recent consensus regarding their phylogenetic relations (Wolfe et al., 2019). Blue boxes indicate translated and smaller grey ones untranslated sequences while lines represent introns. Numbers above the exons and those in italics below the introns indicate their sizes in nucleotides. Where there are two numbers separated by a slash for the two *Penaeus* sequences, the first refers to *P. vannamei* and the second to *P. monodon.* In the case of *Portunus* the + sign is used to separate the nucleotides for the untranslated 5’ and 3’ prime sequences from those of the coding sequences. The sequences for *Penaeus vannamei* and *P. monodon, Portunus trituberculatus, Procambarus virginalis* and *Palaemon carinicauda* are derived from their genome assemblies, the *Cherax quadricarinatus* genome sequence was corrected with reads from genome SRAs and the exon-intron boundaries of the genes from *Stenopus hispidus, Procambarus clarkii, Coenobita clypeatus* and *Eriocheir sinensis* were obtained using Trinity on genome SRAs from these species. The exon sizes of *P. clarkii* and *P. virginalis* are identical, the intron sizes indicated are from the *P. viriginalis* assembly. Question marks indicate exons that are likely present (*Coenobita*) or are present but could not be completely identified (*Eriocheir*).

Trinity does not produce complete gonadulin 2 transcripts for *Eriocheir* or *Carcinus* gonadulin 2; both these sequences, like their *Portunus* ortholog, contain repetitive stretches, making assembly of a transcript difficult. It was also imposssible to find the *Eriocheir* gonadulin 2 gene in either of the two genome assemblies. Genomic SRAs of this species show two very similar, but significantly different genomic sequences for what is likely the first exon of this gene in addition to the repetive sequence. It can therefore not be excluded that this gene is evolving into or has already become a pseudogene. Alternatively, it can neither be excluded that in this species the second gonadulin gene has been duplicated. It is worth noting that all gonadulin gene introns are phase 1, which facilitates the exon shuffling that appears to have occurred with these genes (Fig. 4).

### Receptors

Neuropeptides and hormones need receptors to exert their physiological activity. Insulin-related peptides are somewhat unusual as they act on two different types of receptors, receptor tyrosine kinases (RTKs) and leucine-rich repeat G-protein coupled receptors (LGRs). It has previously been reported that there are four different types of decapod RTKs (Herran et al., 2018) and this was confirmed here for a number of species. Although virtually complete RTK sequences could either be recovered from the databases or using Trinity on a collection of transcriptome SRAs, others were only partially identified. These sequences were combined with RTKs previously identified from *Macrobrachium rosenbergii, Sagmariasus verrauxi* and *Fenneropenaeus sinensis* (Sharabi et al., 2016; Aizen et al., 2016; Guo et al., 2018) and four RTKs from the cockroach *Periplaneta americana,* identified here, for the construction of a phylogenetic tree. The results confirms the existence of four different decapod RTKs and shows that decapods and insects independently evolved four paralogs (Fig. 5; Fig. S2).

**Fig. 5.**
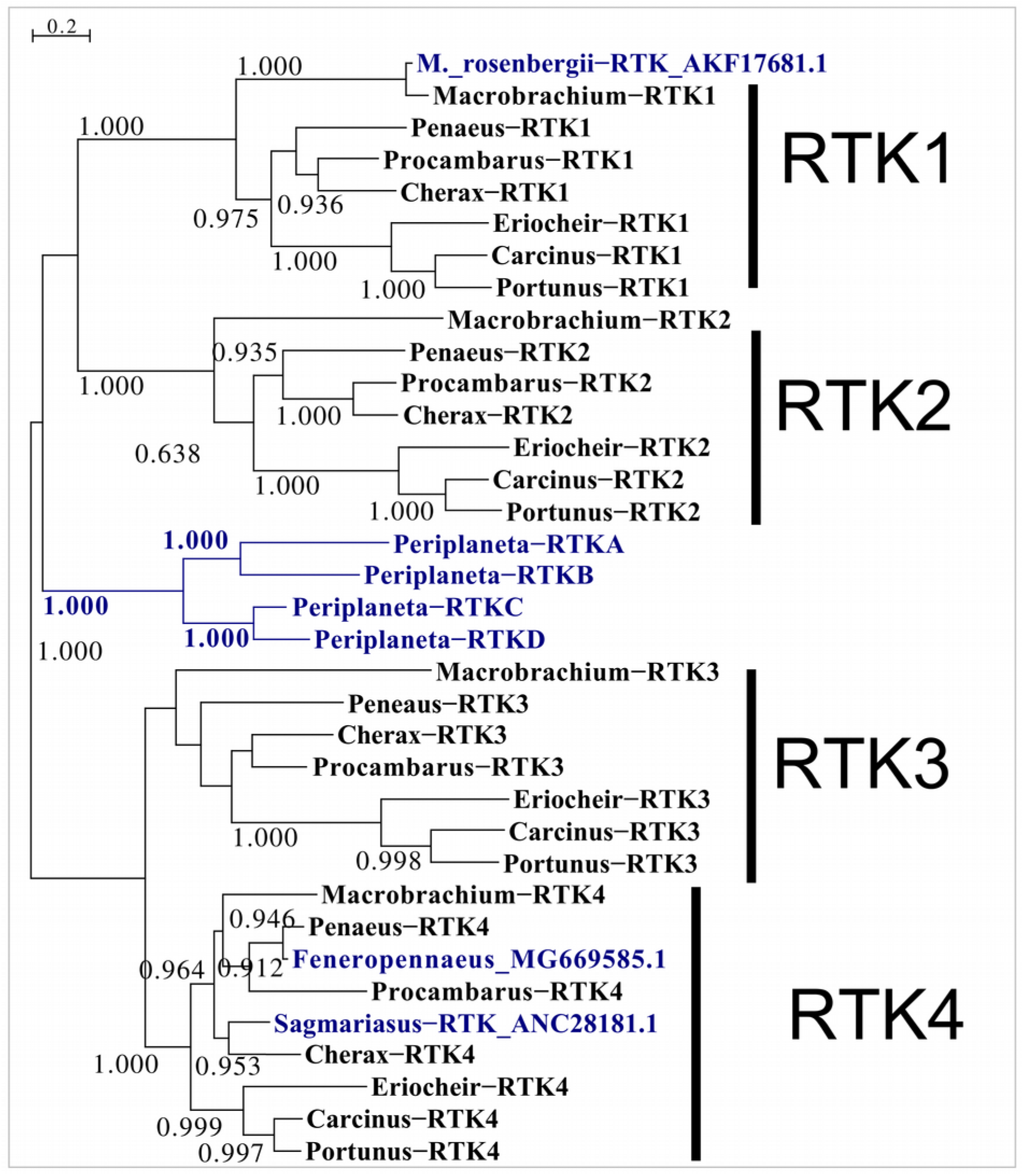
Phylogenetic tree of various decapod RTKs. For comparison sequences of the four *Periplaneta* RTKs have been added, as well as those of three previously identified decapod RTKs, for two of those, the ones from *Fenneropenaeus* and *Sagmariasus*, there is evidence that they interact with the androgenic insulin-like peptide. Sequences are provided in spreadsheet1.

Two arthropod LGRs have either been shown or suggested to be receptors for insulin-related peptides. LGR3 is the receptor for dilp 8 (Vallejo et al., 2015; Gontijo and Garelli, 2018), while LGR4 has been suggested to be the receptor for relaxins, the dilp 7 orthologs (Veenstra, 2014). Many arthropods, but not holometabolous insect species, have another LGR, orthologs of GRL101 from the pond snail *Lymnaea stagnalis* (Tensen et al., 1994). These LGRs are closely related to LGR3 and LGR4 and generally slightly more similar to LGR4 than to LGR3. It suggests, that like LGR3 and LGR4, they might also have an insulin-like ligand. For this reason these receptors were also included and called here LGR5s. Decapods have typically one LGR4, and two LGR3s. The number of LGR5s are not clear, but there often are at least three such receptors. A phylogenetic tree of the transmembrane regions of these receptors and their orthologs from *Periplaneta* and *Drosophila* illustrates their occurrence in a number of decapod species (Fig 6) and their phylogenetic relations.

**Fig. 6.**
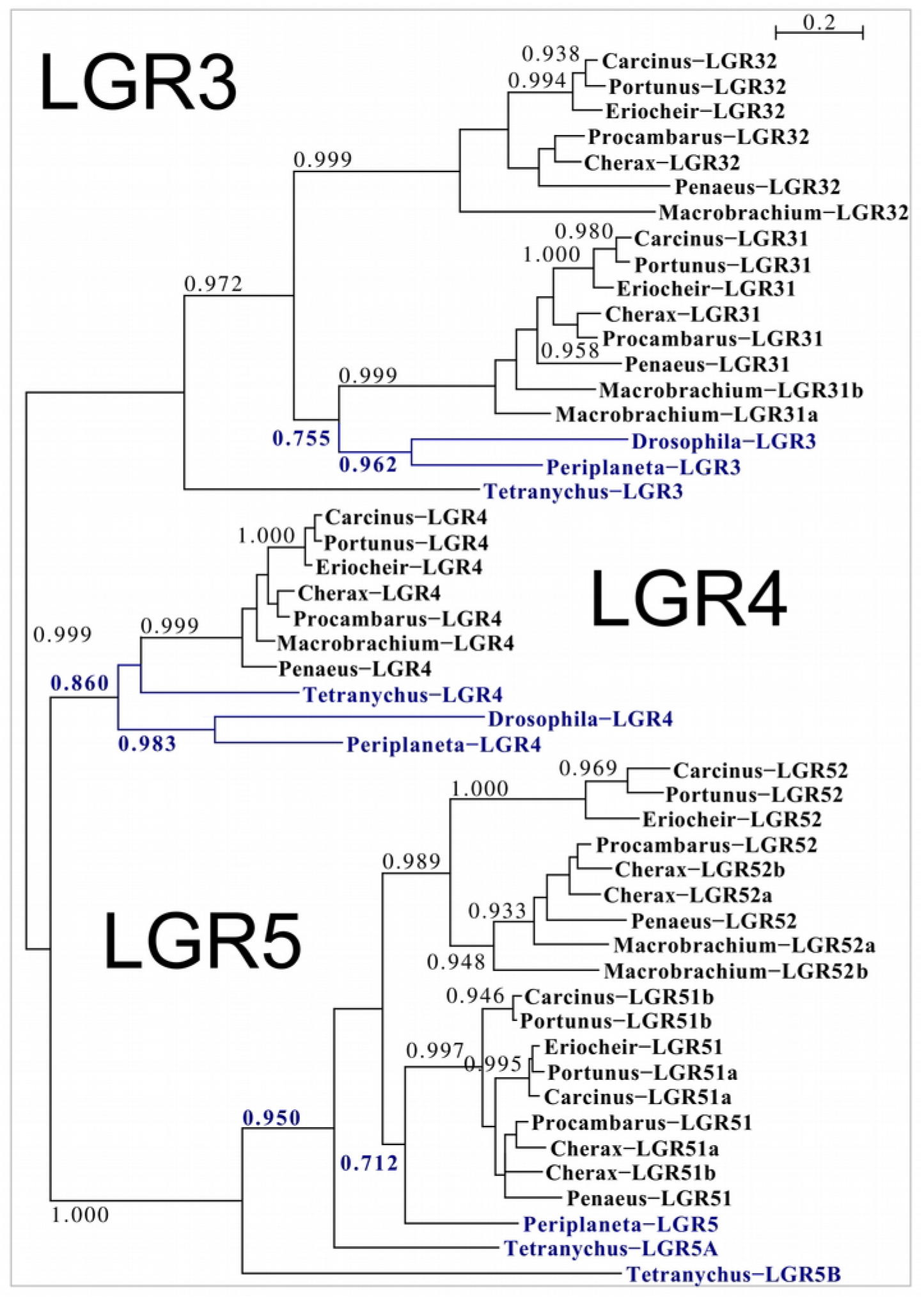
Phylogenetic tree based exclusively on the transmembrane regions of various decapod LGRs that might function as receptors for insulin-related peptides. For comparison sequences of orthologous receptors from the spider mite *Tetranychus urticae*, the fruit fly *Drosophila melanogaster* and the cockroach *Periplaneta americana* have been added. The only deorphanized receptor is *Drosophila* LGR3 which has *Drosophila* ilp 8 for ligand, while the LGR4 are likely ligands for relaxins. Note that *Drosophila* LGR3 has two decapod orthologs, while *Drosophila* LGR4 has only one. On the other hand decapods have several orthologs of *Periplaneta* LGR5. Sequences are provided in spreadsheet1.

### Expression

An attempt has been made to look at expression of the various insulin-related peptides and their putative receptors using the large amount of transcriptomic SRAs available. This approach is obviously limited by a number of factors. Notably, these experiments were done by a large variety of laboratories using different species, different tissues and different physiological conditions. There is a lot of data (spreadsheet2) of which here only some of the more salient results are summarized.

Although the number of samples for the two *Penaeus* species is limited, gonadulin expression in the eyestalk, ovary and the testis of of this species appears to be low. In other decapods gonadulin 1 RNAseq reads are most abundant in the eyestalk as well as the testis, but not in the ovary. *Macrobrachium* appears to be an exception as in this species gonadulin 1 is also expressed in the ovary. Expression of gonadulin 1 is more widespread than that of gonadulin 2, which seems often more limited to the ovary. Nevertheless, in *Procambarus*, the only species where such tissues have been analyzed, it is also expressed in hematopoetic tissue and the anterior proliferation center of the brain; a very low level of expression is also observed in the brain of *Portunus*. The expression of gonadulin 3, a peptide that has only been found in *Macrobrachium,* was only observed in the ovary. Most intriguing is the very variable level of expression of the gonadulin genes in the gonads. This is prominently illustrated by the number of gonadulin RNAseq reads in a number of *Portunus* SRA experiments (Table 1). This highlights a general problem in using SRAs; there there are very few studies that compare different physiological conditions in the same tissue and hence it is very difficult to know why some SRA experiments yield large read numbers for some peptides and others few or none. That the physiological state is important is nicely illustrated by the expression of relaxin in the ovary of *Penaeus monodon*, where the expression of relaxin seems to be correlated with that of vitellogenin (Table S7).

**Table 1.**
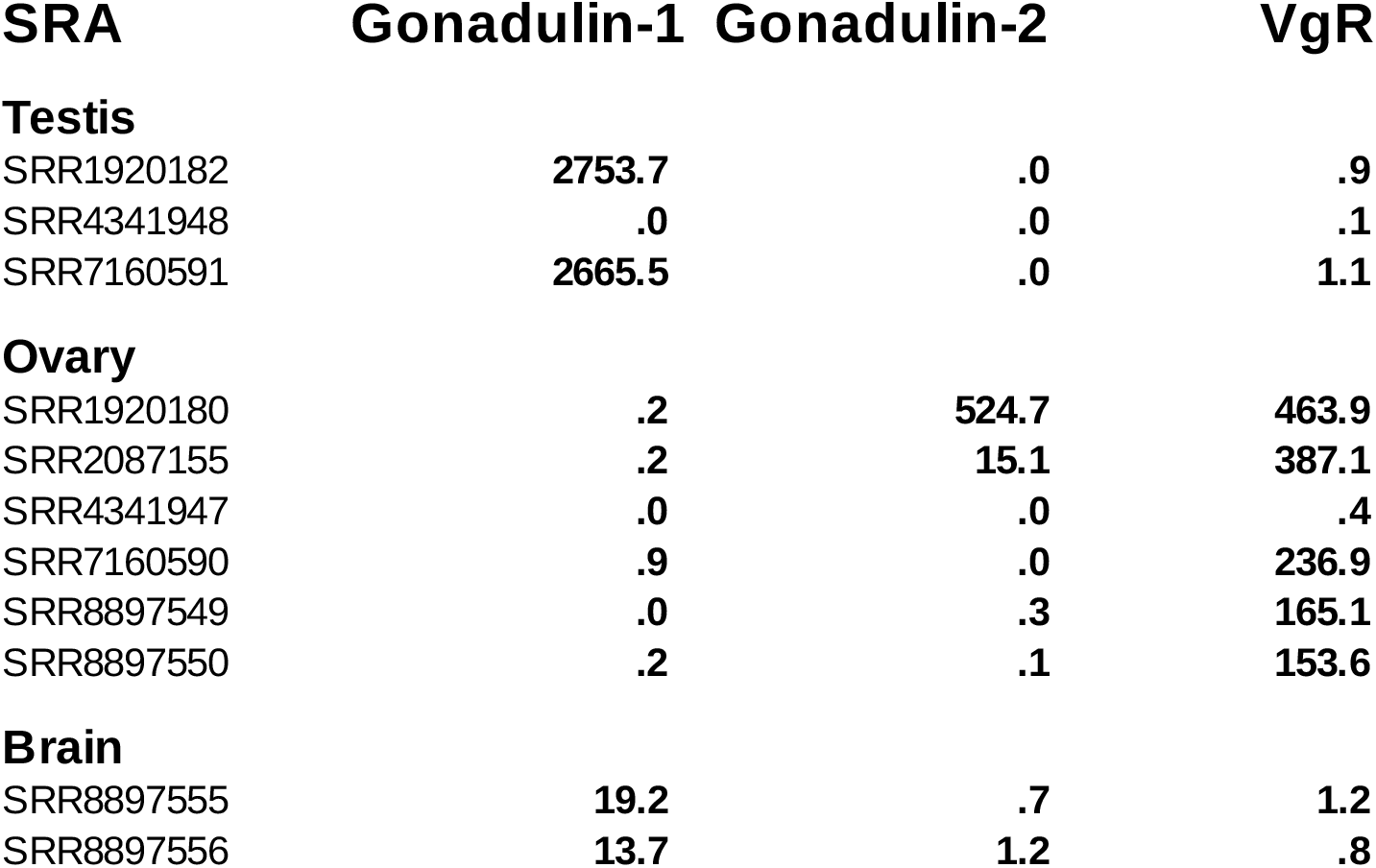
Expression of gonadulin 1 and −2 in the testis, ovary and brain of *Portunus trituberculatus*. Numbers are the number of RNAseq reads per million spots in the various SRAs. Note that expression of gonadulin 2 by the ovary is extremely variable, varying more than a thousand-fold in these samples. Such variation of expression is also present in the testis SRAs. VgR stands for vitellogenin receptor, as the SRA SRR1920180 and SRR1920182 have been mixed up and the expression of the vitellogenin receptor allows unambiguous identification of the ovary SRA. For more extensive data see spreadsheet2.

The RNAseq data suggest expression of androgenin not only in the androgenic gland, but also in the hepatopancreas and/or eyestalk in various species analyzed here. Similar findings have been reported previously (*e.g.* Chung, 2014; Huang et al., 2014; Levy et al., 2017; Shi et al., 2019). Expression levels of both insulin and relaxin are usually low, with insulin mostly expressed in the eyestalk (and nervous system) and in *Procambarus* and *Portunu*s also in the hepatopancreas. However, in *Penaeus vannamei* insulin is expressed in both the testis and the ovary, while in *P. monodon* the ovary expresses relaxin but the the transcriptome SRAs provide no evidence for the expression of insulin in either the ovary or the testis. There are generally very few relaxin RNAseq reads with most of it in the eyestalk and the nervous system, while in *Penaeus modon,* but not *P. vannamei,* there are large numbers in some ovary samples. One hepatopancreas SRA (SRR10846612) out of six from the same *M. nipponense* experiment has a very large number of relaxin reads, suggesting that this particular sample may have been contaminated with another tissue.

It has been reported that in the penaeid *Fenneropenaeus* the RTK4 ortholog is exclusively expressed in the testis (Guo et al., 2018). In the SRAs analyzed here reads for RTK4 orthologs are found mostly, but not exclusively in the testes and androgenic glands. In *Penaeus monodon* RTK3 is also almost exclusively expressed in the testis, unfortunately there is insufficient data for *P. vannamei*, but in other species, RTK3 seems to be more widely expressed. Interestingly, RTK2 orthologs are especially abundant in ovary SRAs, but it is also expressed in many other tissues, including the androgenic gland where at least in some species this RTK seems as highly expressed as in the ovary (Suppl. Data, spreadsheet2).

Within a single species the various LGR are often expressed more strongly in some tissues, but a pattern that was reproduced between the different species was not found (spreadsheet2).

## Discussion

Three different types of insulin-related peptides have previously been shown to be commonly present in decapod crustaceans, the relatively well known androgenin, relaxin, and a third peptide, insulin, that shares characteristics with its mammalian homonyme. The similarity of these peptides with their putative orthologs from insects suggested that the androgenin might be an ortholog of dilp 8 (Veenstra, 2016). However, this hypothesis was challenged by the description of yet another type of decapod insulin-like peptide by Chandler et al. (2017). I show here that this novel type of insect-related peptides is ubiquitously present in these crustaceans and propose to call them gonadulins, as they are commonly expressed by the gonads, sometimes in very large quantities.

Although it remains possible that decapods still have other insulin-like peptides, it is interesting to note that no C-terminally extended insulin was found. Such C-terminally extended insulins are present in several insect species and have sometimes been called insulin-like growth factors as they share characteristics with their homonymes (*e.g.* Defferrari, et al., 2016; Okada et al., 2019).

In several species only incomplete transcripts were obtained for one of the gonadulin genes, but it is nevertheless likely that most decapod species have two such genes. However, several arguments can be advanced to suggest that the penaeids and perhaps all Dendrobranchiata, have only one gonadulin gene. Thus evidence for a second gene was not found in the large number of transcriptome SRAs for these species, nor could such a gene be found in the genome assemblies of two *Penaeus* species or the genome SRA for *Fantapenaeus duorarum*. Interestingly, the *Penaeus* gene has three coding exons and the two introns between them are phase 1 (Fig. 4). Both deletion or duplication of the second coding exon would thus keep the reading frame intact. Deletion of this exon yields a gene structure as seen in the gonadulin 2 and 3 genes, while its duplication would produce one similar to the gonadulin 1 genes of *Stenopus, Procambarus* and *Cherax*. This suggests that a gene duplication may have occurred after the Dendrobranchiata diverged from the other decapods. In a genomic SRA for *Stenopus hispidus* a second gonadulin gene could not be found, although this is not definitive proof that this species lacks a second gene, given the 13-fold coverage of its genome, it is a distinct possibility. Finally, at least in some palaemonidae there are three gonadulin genes, as shown by the transcriptome data from two *Macrobrachium* species and the *Palaemon carinicauda* draft genome.

There are thus (at least) four different types of insulin-like peptides in decapods, *i.e.* relaxin, androgenin, gonadulin and insulin. There are also four different types of insulin RTKs, but it is unknown whether each of the four insulin-like peptides has its own RTK or whether one or more insulin-related peptides acts on more than one RTK. Of course, it also remains possible that one or more of these RTKs use neither of these peptides, but yet another ligand. Within this context it is interesting that orthologs of what has been labeled RTK4 here have been shown to interact with androgenin in two different species (Aizen et al., 2016; Guo et al., 2018). In *Fenneropenaeus chinensis* RTK4 is exclusively expressed in males and the expression data analyzed here for the closely related *P. vannamei* show that this receptor is indeed mainly expressed in testes. However, in other species RTK4 orthologs are more widely expressed. Although RTK2 seems to have its highest expression in the ovary, its tissue specificity is clearly not limited to females, as it is also abundantly expressed in other tissues, including the androgenic gland.

There are also several different LGRs that might be activated by these insulin-like peptides. These can be differentiated in three different main types, LGR3, LGR4 and LGR5, and even more subtypes. The *Drosophila* LGR3 ortholog has been shown to be activated by dilp 8 (Vallejo et al., 2015; Gontijo and Garelli, 2018) and one would expect that the decapod LGR3s would have a similar ligand. Although I previously suggested that this might be androgenin, it now seems much more likely that this would be one or more of the gonadulins, that have a primary amino acid sequence that is more similar to that of *Drosophila* ilp 8 (Fig. S1).

*A priori* one or more of the RTKs and LGR3 would seem to be the most likely receptors for gonadulins, given its identity as an insulin-like peptide (RTK) and its structure that is somewhat similar to dilp 8 (LGR). One of the intriguing characteristics of insulin-related peptides is that they act both through LGRs and RTKs, but to my knowledge there has so far not been an example where a single peptide acts on receptors of both types.

Decapods have only one LGR4, this receptor is present in the same genomes, from both deuterostomes and protostomes, that also have an ortholog of the relaxin gene, and it is absent from genomes that lack a relaxin gene. This strongly suggests that relaxin may be its ligand (*e.g.* Veenstra, 2014, 2019). Complete relaxin transcripts are so far lacking from several decapods although individual reads containing partial relaxin sequences have been found in several cases (Veenstra, 2016). This suggests that the absence of complete transcripts does not reflect an absence of the gene, but rather the lack of transcript analysis from the tissue where the gene is expressed. If the major site of relaxin expression in decapods would be similar to that in *Drosophila* (Yang et al., 2008; Miguel-Aliaga et al., 2008), one would expect relaxin producing neurons in the abdominal ganglia. As this tissue is rarely, if ever, used for RNAseq analysis in decapods, the scarcity of complete relaxin transcripts is perhaps not surprising. Although relaxin reads are occasionally found in hepatopancreas SRAs, these could be due to the accidental inclusion of (a part of) an abdominal ganglion. The abundant amount of relaxin reads in a single SRA *Macrobrachium nipponense* hepatopancreas might represent such an accident.

The large amount of transcriptome SRAs identify tissues where gonadulins are expressed, it also reveals that expression of the decapod insulin-related peptides is rather variable. Thus, whereas the the ovary of *Penaeus monodon* reveals significant relaxin expression, in *P. vannamei* it makes insulin. It is presently impossible to know whether these apparent differences are authentic, or whether they reflect different physiological states of the animals when they were sacrificied. The extremely variable expression of gonadulins by the gonads of *Portunus trituberculatus,* where expression levels vary more than thousand-fold, is also very intriguing and reinforces the conclusion that it is impossible to draw definitive conclusions as to the expression of these peptides. The expression of one of the gonadulins in hematopoetic tissue and the anterior proliferation center of the brain in *Procambarus clarkii* suggest that it, like other arthropod insulin-related peptides, could be a kind of growth hormone, but experiments will be needed to elucidate their function.

## Acknowledgements

I thank three anonymous reviewers for their constructive criticism and help for improving this manuscript. As always for this type of research I am analyzing data I did not collect myself and using methods developed by others. Without either these tools or data, I would not be able to do this and I express my sincere gratitude to all of those who made this possible. I apologize to all those whose SRA data I used without citing the papers that described them; there are simply too many. Institutional funding from the CNRS is also acknowledged.

## Notes

### Competing Interest Statement

The authors have declared no competing interest.

### Summary of Updates

This version of the manuscript has been revised to update the following: Clarify text, correct spelling errors, improve Figs 1 and 4 and add an addition figure New Suppl. Fig. 1 to more clearly demonstrate the existence of four types decapod insulins.

