## SupplementaryData for "Gonadulins, the fourth type of Insulin-related peptides in Decapods"

by Jan A. Veenstra

### Contents

The supplementary data of this manuscript consist of three documents, this pdf file and two spreadsheets. Spreadsheet-1 contains the protein and coding sequences of insulin-like peptides and their putative receptors from eight species, *i.e. Carcinus maenas, Cherax quadricarinatus, Eriocheir sinensis, Macrobrachium nipponense, Penaeus monodon, Penaeus vannamei, Portunus trituberculatus* and *Procambarus clarkii.* Please note that not all these sequences are complete and the larger ones are likely to contain errors. Spreadsheet-2 shows the presence of reads corresponding to each of these proteins in a number of SRAs of the same eight species. These numbers are expressed as both the total amount of half-reads in each SRA, in blue, and their relative numbers, expressed as per million spots, in bold black.


Figure S1. Phylogenetic tree of decapod insulin-related peptides page 2
Figure S2. Phylogenetic tree of decapod RTKs based on the tyrosine kinase domain page 3
Table S1. Transcriptome SRAs used page 4
Table S2. Decapod relaxin precursors page 6
Table S3. Decapod gonadulin 1 precursors page 8
Table S4. Decapod gonadulin 2 precursors page 10
Table S5. Decapod gonadulin 3 precursors page 12
Table S6. Decapod insulin precursors page 12
Table S7. Vitellogenin and relaxin expression in the ovary of Penaeus monodon page 15


Fig. S1. Phylogenetic tree of decapod insulin-related precursors. This tree is not a true phylogenetic tree, but uses the methods employed to make such a tree. It is best considered a similarity tree, as it clusters similar sequences together. It is evident that there are four different types of decapod insulin precursors, but the branch probabilities (only the interesting ones are indicated) are far too small to make any conclusions as to putative phylogenetic relationships. Note though that *Drosophila* ilp 7 is very similar to the decapod relaxins, and that *Drosophila* 8 clusters with the gonadulins and *Drosophila* ilps 1-5 with the insulins, albeit it with a very low branch probability.

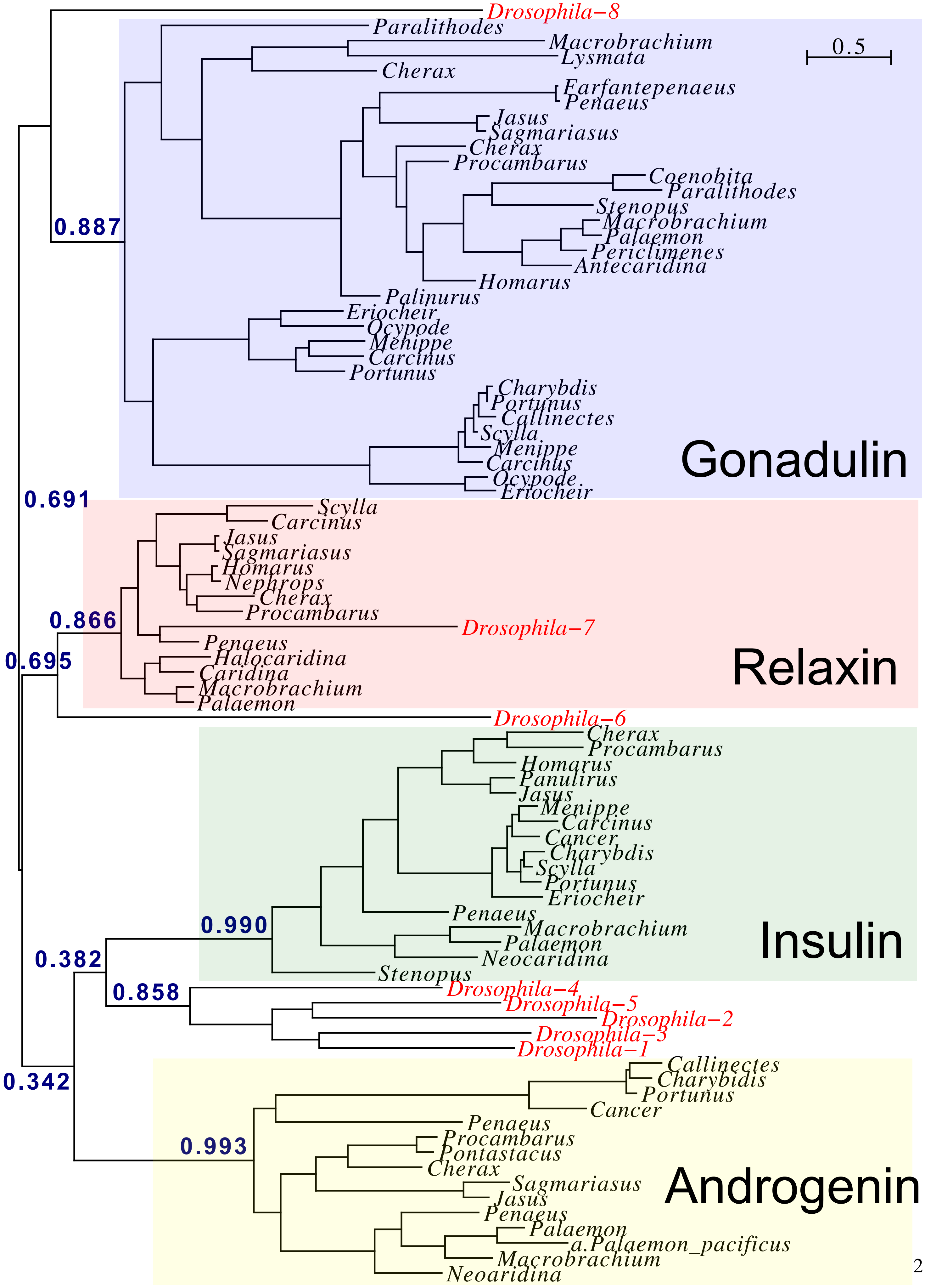

Drosophila−8
Paralithodes
Macrobrachium
Lysmata
Cherax
0.5
Farfantepenaeus
Penaeus
Jasus
Sagmariasus
Cherax
Procambarus
Coenobita
Paralithodes
Stenopus
Macrobrachium
Palaemon
Periclimenes
Antecaridina
Homarus
Palinurus
Eriocheir
Ocypode
Menippe
Carcinus
Portunus
Charybdis
Portunus
Callinectes
Scylla
Menippe
Carcinus
Ocypode
Eriocheir
Gonadulin
0.887
0.691
Scylla
Carcinus
Jasus
Sagmariasus
Homarus
Nephrops
Cherax
Procambarus
Drosophila−7
Penaeus
Halocaridina
Caridina
Macrobrachium
Palaemon
Relaxin
0.866
0.695
Drosophila−6
Cherax
Procambarus
Homarus
Panulirus
Jasus
Menippe
Carcinus
Cancer
Charybdis
Scylla
Portunus
Eriocheir
Penaeus
Macrobrachium
Palaemon
Neocaridina
Stenopus
Insulin
0.990
0.382
0.858
Drosophila−4
Drosophila−5
Drosophila−2
Drosophila−3
Drosophila−1
0.342
Callinectes
Charybidis
Portunus
Cancer
Penaeus
Procambarus
Pontastacus
Cherax
Sagmariasus
Jasus
Penaeus
Palaemon
a.Palaemon_pacificus
Macrobrachium
Neoaridina
0.993
Androgenin

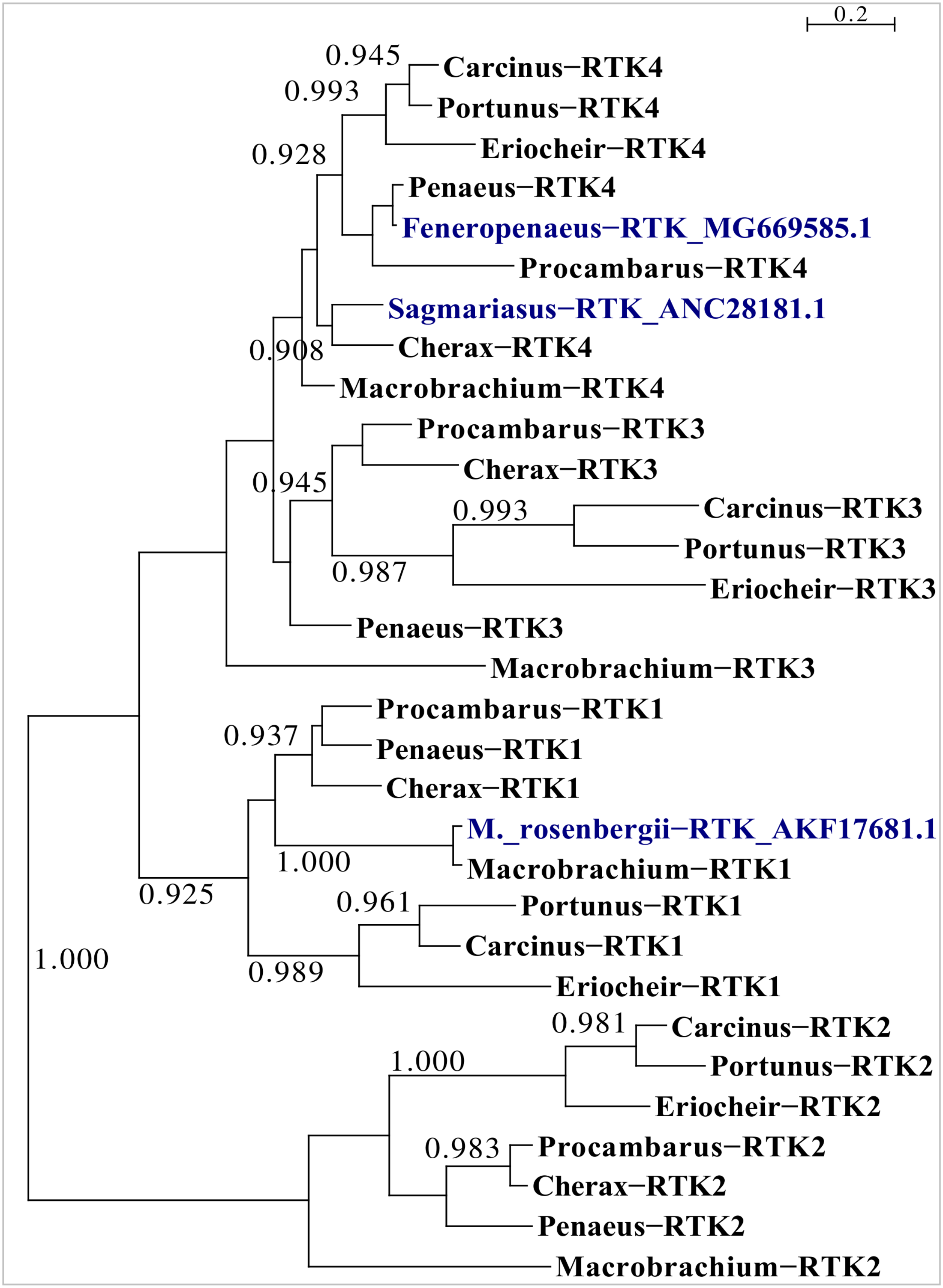


**Figure S2.** Phylogenetic of various decapod RTKs. For comparison sequences of three previously identified decapod RTKs have also been included. For two of those, the ones from *Fenneropenaeus* and *Sagmariasus*, there is evidence that they interact with the androgenic insulin-like peptide.

**Table S1.** Transcriptome SRAs used.

***Carcinus maenas***: SRR1564428, SRR1572181, SRR1586326, SRR1589617, SRR1612556, SRR1632279, SRR1632285, SRR1632289, SRR1632290, SRR1632291, SRR1632292, SRR1632293, SRR6169949, SRR6169951, SRR6169952, SRR6169953, SRR6169954, SRR6169955, SRR6169956, SRR6169957, SRR6169958, SRR6169959, SRR6169960, SRR6169961, SRR6169963, SRR6169964, SRR6169965, SRR6169966, SRR6169967, SRR6169968, SRR6169971, SRR6169987, SRR6169990, SRR6169991, SRR6169996, SRR6169997, SRR6169998, SRR6169999, SRR6170000, SRR6170001, SRR6170002, SRR6170003, SRR6169950, SRR6169969, SRR6169970, SRR6169962, SRR6169992, SRR6169993, SRR6169994, SRR6169995, SRR6169988, SRR6169989, SRR6169972, SRR6169973, SRR6169974, SRR6169975, SRR6169976, SRR6169977, SRR6169978, SRR6169979, SRR6169980, SRR6169981, SRR6169982, SRR6169983, SRR6169984, SRR6169985 and SRR6169986.

***Cherax quadricarinatus***: ERR391748, ERR391749, ERR391750, ERR391751, ERR391752, SRR1814154, SRR4417033, SRR4417034, SRR9903454, SRR9903455, SRR9903456, SRR9903457, SRR9903458, SRR9903459, SRR10023618, SRR10023619, SRR10023621, SRR10023622, SRR10023623, SRR10023624, SRR10023625, SRR10023626, SRR10023627, SRR10023628, SRR10023629, SRR10023630, SRR10023632, SRR10023633, SRR10023634, SRR10023635, SRR10023636, SRR10023637, SRR10023638, SRR10023639, SRR10023640, SRR10023641, SRR10023643 and SRR10023644.

***Eriocheir sinensis***: ERR336998, SRR546086, SRR579531, SRR579532, SRR1555734, SRR1576649, SRR2073826, SRR2170964, SRR2170970, SRR2180019, SRR2180020, SRR3056867, SRR3056869, SRR3623083, SRR3623084, SRR3623085, SRR3623086, SRR3623087, SRR3623088, SRR3623089, SRR3623090, SRR5054211, SRR5054212, SRR5054223, SRR5169063, SRR5169066, SRR5630805, SRR5630899, SRR5894901, SRR5895954, SRR7052450, SRR7052455, SRR7345569, SRR7345578, SRR7345579, SRR7530968, SRR7530969, SRR7585361, SRR7777398, SRR7777399, SRR7777400, SRR7777401, SRR7777402, SRR7777403, SRR7777404, SRR7777405, SRR7777406, SRR7777407, SRR7777408, SRR7777409, SRR7905033, SRR7905034, SRR7905035, SRR7905036, SRR7984118, SRR7984119, SRR7984120, SRR8135129, SRR8135130, SRR8135131, SRR8290193, SRR8290194, SRR8290195, SRR8290196, SRR8290197, SRR8290198, SRR8616629, SRR8618110, SRR7905034, SRR8631920, SRR9179340, SRR9179341, SRR9179342, SRR9179343, SRR9179344, SRR9179345, SRR9179346, SRR9179347, SRR9179348, SRR9179349, SRR9179350, SRR9179351, SRR9663142, SRR9663143, SRR9663144, SRR9663145, SRR9673747, SRR9673748, SRR9673749, SRR9673750, SRR9673751, SRR9673752, SRR9673753, SRR9673754, SRR9673755, SRR9822782, SRR9822783, SRR9822784, SRR9964280, SRR9964281, SRR9964282, SRR10753369, SRR10753370, SRR10753372, SRR10753373, SRR10753378 and SRR10753379.

***Macrobrachim nipponense***: SRR2362707, SRR2363138, SRR3196791, SRR3196792, SRR3709263, SRR4066756, SRR4066757, SRR4066758, SRR4066759, SRR4066760, SRR4066761, SRR4066762, SRR4066763, SRR4066764, SRR4066765, SRR4066766, SRR4066767, SRR4292179, SRR5124602, SRR5124603, SRR5124604, SRR5124605, SRR5124606, SRR5124607, SRR5229543, SRR5229544, SRR5229545, SRR5229546, SRR5229547, SRR5229548, SRR5229549, SRR5229550, SRR5709655, SRR5709656, SRR5709657, SRR5709658, SRR5709659, SRR5709660, SRR5709661, SRR5709662, SRR5709663, SRR5709664, SRR5709665, SRR5709666, SRR5787977, SRR5787978, SRR5787979, SRR5787980, SRR5787981, SRR5787982, SRR5787983, SRR5787984, SRR5787985, SRR5787986, SRR5787987, SRR6820538, SRR6820539, SRR6820540, SRR6820541, SRR6820542, SRR6820543, SRR6820544, SRR6820545, SRR6820546, SRR6986421, SRR6986422, SRR6986423, SRR6986424, SRR6986425, SRR6986426, SRR6986427, SRR6986428, SRR6986429, SRR6986430, SRR6986431, SRR6986432, SRR6986433, SRR6986434, SRR6986435, SRR6986436, SRR6986437, SRR6986438, SRR6986439, SRR6986440, SRR6986441, SRR6986442, SRR6986443, SRR6986444, SRR7250810,

SRR7250811, SRR7250812, SRR7250813, SRR7250814, SRR7665577, SRR7665578, SRR7665579, SRR7665580, SRR7665581, SRR7665582, SRR7665583, SRR7665584, SRR7665585, SRR8421823, SRR8421824, SRR8421825, SRR8421826, SRR8421827, SRR8421828, SRR8421829, SRR8421830, SRR8421831, SRR8421832, SRR8421833, SRR8421834, SRR8422891, SRR8422892, SRR8422893, SRR8422894, SRR8422895, SRR8422896, SRR8422897, SRR8422898, SRR8422899, SRR8422900, SRR8422901, SRR8422902, SRR8422903, SRR8422904, SRR8422905, SRR8422906, SRR8422907, SRR8422908, SRR8989425, SRR8989426, SRR8989427, SRR8989428, SRR8989429, SRR8989430, SRR8989431, SRR8989432, SRR8989433, SRR8989434, SRR8989435, SRR8989436, SRR8989437, SRR8989438, SRR8989439, SRR8989440, SRR9333612, SRR9333613, SRR9333614, SRR9333615, SRR9333616, SRR9333617, SRR9333618, SRR9333619, SRR9333620, SRR9333621, SRR9333622, SRR9333623, SRR9333624, SRR9333625, SRR9333626, SRR10824559, SRR10824560, SRR10824561, SRR10846612, SRR10846613 and SRR10846615.

***Macrobrachium rosenbergii***: DRR023219, DRR023253, SRR345608, SRR345609, SRR345610, SRR345611, SRR567391, SRR572719, SRR572720, SRR572721, SRR572722, SRR572723, SRR572724, SRR572725, SRR896637, SRR896638, SRR896645, SRR896646, SRR896647, SRR896649, SRR896650, SRR896651, SRR1138560, SRR1138561, SRR1138562, SRR1138563, SRR1138564, SRR1138565, SRR1138572, SRR1138573, SRR1424572, SRR1424574, SRR1424575, SRR1559285, SRR1559286, SRR1559287, SRR1559288, SRR1653452, SRR1653453, SRR1653454, SRR1781508, SRR2082537, SRR2082765, SRR2082766, SRR2082767, SRR2082768, SRR2082769, SRR2082770, SRR3087290, SRR5396640, SRR6179325, SRR6179326, SRR6179327, SRR7938693, SRR7938694, SRR7938695, SRR7938696, SRR7938697, SRR7938698, SRR7942857, SRR7942858, SRR7942859, SRR7942860, SRR7942861, SRR7942862, SRR8426969, SRR8426970, SRR8426971, SRR8426972, SRR8426973, SRR8426974, SRR8432415, SRR8432416, SRR8432417, SRR8432418, SRR8432419, SRR8432420, SRR8432421, SRR8432422, SRR8432423, SRR9670548, SRR9670549, SRR9670550, SRR9670551, SRR9670552, SRR9670553, SRR10158823, SRR10158824, SRR10158825, SRR10158826, SRR10158827 and SRR10158828.

***Palaemon serratus***: SRR8631955, SRR8631956, SRR8631957, SRR8631958, SRR8631959 and SRR8631960.

***Penaeus monodon***: SRR6868116, SRR6868117, SRR6868118, SRR6868119, SRR6868120, SRR6868121, SRR6868122, SRR6868123, SRR6868124, SRR6868125, SRR6868126, SRR6868127, SRR6868128, SRR6868129, SRR6868130, SRR6868131, SRR6868132, SRR6868133, SRR6868134, SRR6868135, SRR6868136, SRR6868137, SRR6868138, SRR6868139, SRR6868140, SRR6868141, SRR6868142, SRR6868143, SRR6868144, SRR6868145, SRR6868146, SRR6868147, SRR6868148, SRR6868149, SRR6868150, SRR6868151, SRR6868152, SRR6868153, SRR6868154, SRR6868155, SRR6868156, SRR6868157, SRR6868158, SRR6868159, SRR6868160, SRR6868161, SRR6868162, SRR6868163, SRR6868164, SRR6868165, SRR6868166, SRR6868167, SRR6868168, SRR6868169, SRR6868170, SRR6868171, SRR6868172, SRR9031899, SRR9031900, SRR9031901, SRR9031902, SRR9031903, SRR9031904, SRR9031905, SRR9031906, SRR9031907 and SRR9031908.

***Penaeus vannamei***: SRR346404, SRR554363, SRR554364, SRR554365, SRR556131, SRR1037362, SRR1037363, SRR1037364, SRR1037365, SRR1037366, SRR1039534, SRR1104080, SRR1104081, SRR1104082, SRR1104083, SRR1104084, SRR1104085, SRR1104086, SRR1104087, SRR1105791, SRR1184416, SRR1407787, SRR1407788, SRR1407789, SRR1407790, SRR1407791, SRR1460493, SRR1460494, SRR1460495, SRR1460504, SRR1460505, SRR1609917, SRR1618514, SRR1951370, SRR1951371, SRR1951372, SRR1951373, SRR2060962, SRR2060963, SRR2060964, SRR2060965, SRR2103851, SRR2103852, SRR2103853, SRR2103854, SRR2103855, SRR2103856, SRR2103857, SRR2103858, SRR2103859, SRR2103860, SRR2103861, SRR2103862, SRR2103863, SRR2103864, SRR2103865, SRR2103866, SRR2895158, SRR3619839, SRR5065708, SRR5065709, SRR5065710, SRR5065711, SRR5065712, SRR5065713, SRR5065714, SRR5065715, SRR5065716, SRR5134182,

SRR6147939, SRR6147940, SRR6147941, SRR6147942, SRR6147943, SRR6147944, SRR6466294, SRR6466295, SRR6466296, SRR6466297, SRR6466298, SRR6466299, SRR6466300, SRR6466301, SRR6466302, SRR6466303, SRR6466304, SRR6466305, SRR6466306, SRR6466307, SRR6466308, SRR6466309, SRR6466310, SRR6466311, SRR6466312, SRR6466313, SRR6466314, SRR6466315, SRR6466316, SRR6466317, SRR6466318, SRR6466319, SRR6466320, SRR6466321, SRR6466322, SRR6466323, SRR6466324, SRR6466325, SRR6466326, SRR6466327, SRR6466328, SRR6466329, SRR6466330, SRR6466331, SRR6466332, SRR6466333, SRR6466334, SRR6466335, SRR6466336, SRR6466337, SRR6466338, SRR6466339, SRR6466340, SRR6466341, SRR6466342, SRR6466343, SRR6466344, SRR6466345, SRR6466346, SRR6466347, SRR6466348, SRR6466349, SRR6466350, SRR6466351, SRR6466352, SRR6466353, SRR6466354, SRR6466355, SRR6466356, SRR6466357, SRR6488337, SRR6488338, SRR6488339, SRR6488340, SRR6488341, SRR7453063, SRR7756945, SRR7756946, SRR7756947, SRR7756948, SRR7756949, SRR7756950, SRR7756951, SRR7756952, SRR7756953, SRR7756954, SRR7756955, SRR7756956, SRR7756957, SRR7756958, SRR7756959, SRR7756960, SRR7756961, SRR7756962, SRR7756963, SRR7756964, SRR7756965, SRR7756966, SRR7756967, SRR7756968, SRR7756969, SRR7756970, SRR7756971, SRR7756972, SRR7756973, SRR7756974, SRR7756975, SRR7756976, SRR7756977, SRR7756978, SRR7756979, SRR7756980, SRR7756981, SRR7756982, SRR7756983, SRR7756984, SRR7756985, SRR7756986, SRR7756987, SRR7756988, SRR7756989, SRR7756990, SRR7756991, SRR7756992, SRR7756993, SRR7756994, SRR7756995, SRR7756996, SRR7756997, SRR7756998, SRR7756999, SRR7757000, SRR7757001, SRR7757002, SRR7757003, SRR7757004, SRR7757005, SRR7757006, SRR7986779, SRR9822083, SRR9822084, SRR9822100, SRR9822101, SRR9822102, SRR9822103, SRR9822104, SRR9822105, SRR9822106, SRR9822107, SRR9822108, SRR9822109, SRR9822111, SRR9822112, SRR9822113, SRR9822114, SRR9822115 and SRR9822116.

***Portunus sanguinolentus***: SRR6216182.

***Portunus trituberculatus***: SRR768319, SRR479039, SRR1013694, SRR1013695, SRR1013696, SRR1105793, SRR1168416, SRR1168417, SRR1185310, SRR1630818, SRR1920180, SRR1920182, SRR2087155, SRR4341947, SRR4341948, SRR5807789, SRR5807790, SRR5807791, SRR5807792, SRR5807793, SRR5807794, SRR6233346, SRR6447887, SRR6448507, SRR6448508, SRR6448509, SRR6459731, SRR6678996, SRR6678997, SRR6678998, SRR7160590, SRR7160591, SRR8076048, SRR8076049, SRR8076050, SRR8076051, SRR8133954, SRR8133955, SRR8187066, SRR8187067, SRR8187068, SRR8187137, SRR8393292, SRR8801865, SRR8897547, SRR8897548, SRR8897549, SRR8897550, SRR8897551, SRR8897552, SRR8897553, SRR8897554, SRR8897555, SRR8897556 and SRR9964020.

***Procambarus clarkii***: SRR870673, SRR1144630, SRR1144631, SRR1265966, SRR1509455, SRR1509456, SRR1509457, SRR1509458, SRR3458513, SRR5136645, SRR5136646, SRR5136647, SRR5136648, SRR5136649, SRR5136650, SRR5136651, SRR5136652, SRR5136653, SRR5136654, SRR5136655, SRR5136656, SRR5136657, SRR5136658, SRR5136659, SRR5136660, SRR5136661, SRR5136662, SRR5136663, SRR7601327, SRR8151934, SRR8151935, SRR8156034, SRR10874080, SRR10874081 and SRR10874082.

***Stenopus hispidus***: SRR5140147, SRR10023645, SRR10023646, SRR10023647, SRR10023648, SRR10023649, SRR10023650, SRR10023651, SRR10023652, SRR10023654, SRR10023655, SRR10023656, SRR10023657, SRR10023658, SRR10023659 and SRR10023660.

**Table S2.** Predicted decapod relaxin precursors.

Species name and identifier are indicated; Trinity means that the sequence obtained from Trinity generated transcripts from public SRAs. In yellow the predicted signal peptide, in blue the likely mature peptide that will produced after cleavage by convertase at the proteolytic cleavage sites that are highlighted in black. Cysteine residues have been highlighted in red.

***Carcinus maenas*** GFXF01072889.1, GFXF01126361.1

MFLLMVVVVVVVVDLSTCIDPDLLSEIRNRNARDWQALWSEERLALCRLRLRHNLDAICGKDVYRRSAPPAPPL
QDEDEERKIQEEEEEEEEEEGGVGEVGEKRGLGGGVSLHLPSTRIEINPPPPTPHYKDQGGDSRSPLLSVQQA
NLFVTTWIRDTPPPPKDNFGKPQIDPLRKGGGGSRWHRHPRRSLYYPPHPRHARHAPSITSECCTEVGCTWEEY
AEYCPSSSRLRPGVTLI

***Caridina multidentata*** IABX01041081.1 & IABX01121135.1

---VVLLLAATFTIISSSSAFDQDFLQRIESRTASEWEAAWSEERLALCRARLRHNLDAICGKDIYRR
SSAIQERFKRRAPKCLHRRGLPFNSSGDHNLQRGMRGLNAIVPGPLTAMKIGPSLPDTGQHNEQRKMPLLSVQE
ANLFVTTWVRGQRVTHGRRRRQSQSITSECCTDVGCTWEEYAEYCPISTRARPGVIPI

***Cherax quadricarinatus*** KP006644

MLALTAMFVLGSTSWALESDLIRQIESRTETEWQTLWSKERLSLCRARLRHNLDTICGKDVYRRSLAPPRPAPY
HHIFKRRTDICLQVHDTGGARRVEGEKHLSKSSNRVKRVREVLVNLSPDIIQTSPATDTGQPSVQDRHVHSRYR
SPFLSVHQANLFVTTWVRDHQGRHYRRRRQSSSITAECCTTTGCTWEEYAEYCPTSSRLRAGVALI

***Halocaridina rubra*** GHBK01042169.1

MRRDMVFLLLVATFTFISSSTAFDQEILQRIESSRTASEWEAIWSEERLALCRARLRHNLDAICGKDVYRRSTD
NQGRFKRRPPKCWRGRGLFNGSGKYSPKYERKKSQVSLLSPFAKLNNAQVFSENGQNTKERRSPFLSVQEANLF
VTTWLRGRRLTHGRTRRQSPSITSECCTERGCTWEEYAEYCPTSSRARPGIIPI

***Homarus americanus*** GFUC01099703.1

MVVVIAAILVVVSTSWALEPYLIRQLESRTEAEWEVLWNKERLALCRARLRHNLEAICGKDVYRRSLTPPNHHH
IKRSTDTCLKVHDGDGERDVRDKRAVSVNLPTATIEITPSSPDTGQHNINTRSPFLSVHQANLFVTTWVGGRRG
SHYRRRRQSSSITAECCTTVGCTWEEYAEYCPTSSRLRPGVTPI

***Jasus edwardsii*** GGHM01005531.1,GGHM01014244.1, GGHM01014245.1

MVAADMVVLVLVLAAMLTLVTFSWALDPDLISQIESRTEKEWQELWTEERLTLCRSRLRHNLDAICGKDVYRRS
PMLPPRHRRWSRAKRNTDIFVEVHDTDAVRGDSGKKEKRMRTMSVDLPTTRIEISPSVPDTGQHSTHTRSPFLS
VHQANLFVTTWVGGHHRRRRQSPSITSECCTTVGCTWEEYAEYCPTSSRLRPGVALI

***Macrobrachium nipponense*** GHMG01012391.1

MNKDMVVLVLAATVTLLSSASAFDQSLLQRIESRTASEWQAVWSEERLALCRARLRHNLDAICGKDVYRRSPPG
HQGRYKRRAPKCLRTQAGGTTNNSGDNRSTTTNSNAVMTYPPSAPVVRPSLPDTGQNTDEGRSPFLSVQQANLF
VTTWVRGGGPVHGRRRRQSQSITSECCTAAGCTWEEYAEYCPTSSRVRPGVIPI

***Nephrops norvegicus*** QBX89070.1

MVVVIAAILVVVSTSWALEPDLIRQIGSRTESEWEVLWNKERLALCRTRLRHNLEAICVKDVYRRSLTSPNHHH
IKRSTDICLKVHDSDGEGDIRDKGAVSVNLPTATIEITPSSPDTGQHNIYTRSPFLSVQQANLFVTTWVGGRRG
GHYRRRRQSSSITAECCTTVGCTWEEYAEYCPTSSRLRPGVTPI

***Palaemon serratus*** Trinity

MKKDMVVLVLAATVTLISSTSAFDQNLLQRIDSRTASEWEAAWSEERLALCRARLRHNLDAICGKDVYRRSPGQ
GRQKRRAPKCHRAQGGSNNNDENKTSNFNAVMTYPPVSADVRPSLPDTGQNQEEGRPPFLSVQQANLFVTTWVR
GRSAEHGRRRRQSQSITSECCTAAGCTWEEYAEYCPTSSRVRPGVIPI

***Penaeus monodon*** GGLH01089237.1

MVMSMVLAVFLLCSTSLALDPDFVRQIESRTELEWQTLWSEERLALCRAKLRQNLDAICGKDVFRRSSAGGRRR
DKRDKDRGGRQPLPAESDEAPRVNPSTPDTGQTLDKRRSPFLSVQQANLFVTTWVHDQGGRRRGRSHFRRRRQS
PSITTECCTVAGCTWEEYAEYCPSSNRARLL

***Penaeus vannamei*** GHXV01059825.1
MVMSMMLAVFLLCSTSLALDPDFVRQIESRTELEWQALWSEERLALCRAKLRQNLDAICGKDVFRRSSVERRRR
RDKRDEGRDGSKPLPAESDEVPRANPSTPDTGQAPDKRRSPFLSVQQANLFVTTWVHDQGGRRRGRSHYRRRRQ
SPSITTECCTVAGCTWEEYAEYCPSSNRARFL

***Procambarus clarkii*** GBEV01092320.1
MMALLLAAMFVIAAISWALDPDLIRQIESRTEAEWQTLWSKERLALCRARLRYNLDSICGKDVYRRSLKTPPPS
HHQHQKQHHLVKRTTDICVHVHEAGGESAEDNTEKREKSLDGAESILPSTTIEINPSTPDTGQESVQARSPFLS
VHQANLFVTTWVGGRGRRGPQHRLRRQSPSITAECCTAVGCTWEEYAEYCPTSSRLRAGVTLI

***Sagmariasus verreauxi*** KP006643.1
MLAADMVVLVLAAMLTLVTFSWALEPDLISQIESRTEKEWQELWTEERLTLCRSRLRHNLDAICGKDVYRRSSM
LPPRTRHRRWSRAKRNTDIFLEVHDTDTARGDSRKKEKRMKTMSVDLPTTRIEISPSVPDTGQHSTHTRSPFLS
VHQANLFVTTWVGGHHRHRRQSPSITSECCTTVGCTWEEYAEYCPTSSRLRPGVTLI

***Scylla olivacea*** GDRN01010195.1, GDRN01041251.1
MFVFIVVVMVVVAVNTTTSLDPDLLGEIENRDAHDWEALWSEERLALCRLRLRQDLDSICGKDVFRRSAPPRPA
APPPEPPLYDGDGARKNVEEEEEEEASDECVVGRRGGGRGGRWHRYPSMYPHYPLTPRPPRHAPSITSECCTEV
GCTWEEYAEYCPSSSRLRPGVKLI

### **Table S3.** Predicted decapod gonadulin 1 precursors.

Species name and identifier are indicated; Trinity means that the sequence obtained from Trinity generated transcripts from public SRAs and Artemis that the sequence is predicted from either a draft genome or genomic SRAs. In yellow the predicted signal peptide, in blue the propeptide that will produced after removal of the signal peptide. Possible proteolytic cleavage sites are highlighted in black and cysteine residues in red. See note about processing of the gonadulin precursor in the main text.

***Antecaridina lauensis*** GHBJ01019446.1
MRIIGALALAIVLVAVVETRPPREKRGIKICSARDVKFMATFVCNLHKRSVRSVDAAEFGAHVPGTDGLTALGG
AGHLPRRTICNGGDCGRQIDGVDFHSSDPLNYIQAWLSGNGYPPMMTSGYRNVPARATENGQDMYEENEVGRPN
LLQVLERMGAMRNGIEKRDKEIDWPLVPSMTLGDIRKNCCLRQCRVEDFYGACS

***Callinectes sapidus*** GEID01030804.1
MRVVVGVVAVVALVVAVAEGDSLMMRSKKYKLCTARDINMFVTQLCSGSISRMRVGRHNSHPARGILTIRIPPR
DVRGPVVSPAVIGSAYEFPSYRESLLDTSKYSPSELDTEDEVPAEYYANALTRSKKWLNDQWNAGEARSTMLGV
REKRLTFGDIRQRCCTDGCYASELQSLCA

***Carcinus maenas*** GFXF01213449.1
MRVVVGVVAMVAVVVAVLVAVAEGDSLMMRSKKYKLCTPRDINMLVSQLCSGAISRMRGGRHYSHKAVPGIMTI
RIPPRDVRGPVVSPAKIGSSYEFPSYRESLLETPKLSTSDLDAVDAPYLRPVSYWENQDWDNYKDSSAETASEY
NGNALNPRRRKWWYDQWNAGDPNTAQFVPRMKRTTFGDIRLRCCTEGCYASELQSLCA

***Charybdis feriata*** GGFD01059139.1
---ALVVAVAEGDSLMTRSKKYKLCTARDINMFVTQLCSGAINRMRVGRHNSHPAPGILTIRIPPRGVR
GPVVSPAEIAPYLRPASYWESQGWNNYKESSAETEIPSEYYGNGFTSRSKKWLNDQRNEGEGSSAMIWLRRKRL
TFGDIRQKCCTDGCYASELQSLCA

***Cherax quadricarinatus*** AIU40993.1
MGRCSLRGIGALVFVLATAALLVETRPPYRSRRGMKVCSPRDVKFMATYICNLHRRSVRSVDDFEDDFESPGVS
RLSGVNIPPWRPSRCGNTGRDCGPGLDSSSSLPPLARLHHHSPPAATTITAADFNRWLSLNGYHDLSLQEDSN
GLGNDLVNDPWQAIRENGETNQERENGLVRVNSATHFLGDISLAKRDREVDWPVLVPRSLSDIRRNCCLRECTA
EDFYGACS

***Eriocheir sinensis*** GFBL01084478.1

MRVVVVGVVAVVVLAVAVVGAEVRLTRAKKYRLCTARDVSMLVAELCSGTIGSPMRGGLHRGAGGNAGIMIRSQ
PIAIRGRRRENDGPVFESLPLWENVNNQKYSRDELDRLNGSPDLIPAWDMQDTNSYEDLLAEPGWRVEARPDTF
LSALVRLSRGGERRPGSPLAPGLTHTARDKRQTLSKLRARCCREGCDEAELYTFCV

***Halocaridina rubra*** GHBK01033082.1

MKIVGAFVLAIVLVVVVETRPPREKRGIKICSARDVKFMATFVCNLHKRSVRSVDAEDFGDEVPDTEGVPAFGG
AGHLPRRTICNGEDCRQQIEGNELLSSDPLTYIQAWLAENGYPSAMSSGYRNVPGRATENGHRHLGHYEENGID
RKGLQQVLEHLGVARNAFAKRDKEIDWPVVPSMTLGDIRKNCCLRQCRV

***Homarus americanus*** GEBG01024539.1,GFUC01138962.1,GFDA01112291.1 and GFDA01112292.1

MRATGALVLVMVFLAAMVETHPPRAKRGIKLCSARDVKFMATYVCNLHRRSVRSVDQHSDDNYETTGVEELGGG
NSRAWRSSRCGPSGSDCAPQFDSLPFQPSAVDFGTPAVISMADVERWLSQNGYHRLILPEYSSFPWQTKNGPTN
NGAASNRNIENELYQSNYGTGVMGDLRATMNKRDKEIDFPVMVPRSLSDIRKNCCVRECTAEDFYGACS

***Macrobrachium nipponense*** GHMG01073270.1

MRFIGAFALVVLLVATVAETRPPSPREKRGIKICSARDVKFMATFVCNLHKRSVRSVDLDENEDDDEDFGLPVT
DDDLALLGSLGMVPRRPLCGDDGRDCGRQNKPANITSSQLNYLQAWLEENGLLLSSTLPEERSQTRTSDLRHHL
LRPLLRNIETGSQRGNPRIPLEDLFPSLKGSVTKRDKEIDWPLVPSMTLGDIRKNCCIRQCRVEDFYGACT-

***Macrobrachium rosenbergii*** Trinity

MRFIGAFALVVFLVATVAETRPPSPREKRGIKICSARDVKFMATFVCNLHKRSVRSVDLDENEDDEDFGLPVTD
DDLAMMGSLGMVPRQALCGDNGRDCGRQNKPANITSSQLNYLQAWLEENGLLLPSLPDERSQPRTTNLHHHLLR
PLLRNVETASQRGNPRIPLEDLFPSLKGSVAKRDKEIDWPLVPSMTLGDIRKNCCIRQCRVEDFYGACT

***Macrobrachium tolmerum*** GHDQ01055738.1, GHDQ01055738.1

MRFIGAFALVVFLVATVAETRPPSPREKRGIKICSARDVKFMATFVCNLHKRSVRSVDLGDNDDDDEDFGLPVT
DDDFAMLGSLGMVPRRPLCGDNGRDCGRQIKQANITSSQLNYLQAWLEENGLLLPSLPDERSQPRTSDLRYHLL
RPLLRNVESGSQRGNPRIPLEDLFPSLKGSVAKRDKEIDWPLVPSMTLGDIRKNCCIRQCRVEDFYGACT

***Menippe nodifrons*** from SRR8787123

MRVVMGVVVVVALVVAVAEGDSVMTRTKKYKLCTARDINLFVSQLCSGTLTRMRVGRHSSHPGILTIRVPPRDV
RAPVVSPAEIGSSPEFPSYRESLDSQRYSPSDLDAMNAPYLRPVSYWESQEWNNYKDSSAEPELPSDYYGNDLS
PRSKKWWYDQLSAANPNADMIGLRQRRTRFGDIRQRCCTEGCYASELQSLCVV

***Palaemon carinicauda*** Artemis

MRFVGALALVVVLAAVAETRPPREKRGIKICSARDVKFMATFVCNLHKRSVRSVDLDEEDFALPVSDDDLAMLG
FLGSVPGSVPGRPLCGDNGRDCGRQVKVANITNSQLNYLQAWLAENGLVPSFPNERNQPRMTENHRPLHRNIEY
GLQRETPRALLEDLFPSLKGPVAKRDKEIDWPLVPSMTLGDIRKNCCVRQCRMEDFYGACT

***Palaemon serratus*** Trinity

MRFVGALALVVVLAAVAETRPPREKRGIKICSARDVKFMATFVCNLHKRSVRSVDLDEDDFALPVTDDDLAMLG
FLGSVPGRPLCGENGRDCGRQVKVSNITTSQLNYLQAWLAENGLVPSFPDERSQPRMTENHRPLHRNIEYGLQR
ETPRILLEDLFPSLKGPVAKRDKEVDWPLVPSMTLGDIRKNCCVRQCRMEDFYGACT

***Palaemon varians*** GFPG01044320.1

MRFVGALALVVVLAAVAETRPPREKRGIKICSARDVKFMATFVCNLHKRSVRSVDLDEDDFALPDTDDDLALLG
FLGSVPGRPLCGEDGRDCGRQIKVSNITNSQLNYLQAWLAENGLVPSFPNERNQPRMPDSHRPLHRNIEYGLQR
ENPRILLEDLFPSLKGPVAKRDKEIDWPLVPSMTLGDIRKNCCVRQCRMEDFYGACT

***Paralithodes camtschaticus*** GHJC01030171.1

MRCWVAAAVVVVVCVVGVVVVEPRPQQQHQHYHQEKRGVRLCSARDVKLIATYVCNLHRRSVRSVALDENSEGD
TYIMPGVGGSSVILPWRSLPPCDGPDGDCGGLNSADSPVQQTQRASPPQAPFNMAKRWYALHRFGTAKGMQAEK
ENDITEDYRADSNSIDYAPSLSGALPLSELFPENVSKFIRVNYPKYVTSLLPHKGKRDKEMSLPVVVPRSLSAI
RQDCCVKECNAEDFFGACS

***Penaeus vannamei*** Trinity
MRTLATLVVVALLATASEAGERPSLYKRGQLVHVCTPRDVKMMARFVCSLHRRSVRSSSSSSSSVADSVIVSGP
LFRIPEFYNRPPSRVMKCGENGDDCLPAYDPDYDSASYFLPPSGDLRTNLIKRDKEVVRAVGLTIADVRRKCCL
NGCLPEDFYGACR

***Portunus trituberculatus*** GFFJ01045966.1
MRVVVGVVAVVALVVAVAEGDSLMTRSKKYKLCTARDINMFVTQLCSGALTRMRVGRHNSHPAPGILTIRIPPR
GVRGPVVSPAEIGSSYEFPSYRESLLDTQNYSPSELDTVDAPYLRPVSYWESQSWNNYKESSAETEVPSEYYGN
GLTSRSKKWLNDQRNEGEGSSAMIWLRRKRLTFGDIRQKCCTDGCYASELQSLCA

***Procambarus clarkii*** GBEV01005034.1, GARH01003670.1
MRGVGALMLVAVLTAATLVVETRPPPRNKRGIRLCSARDVKIMATYVCNLHRRSVRSVHEPDNTFRSSDAASLV
DVSNRPWRSSPCGNTGWDCIPEFDSHPIQSSVVDVDDAASITLADIQRWLSANGYHRMFLPENSNNLWPAKIGM
INENADINQDTENGFGRVNYAIPFLGDLRTNMAKRDKELDWPVVAPRSLGDVRKNCCLRECSVEDFYGACS

***Procambarus virginalis*** Artemis
MRGVGALMLVAVLTAATLVVETRPPPRNKRGIRLCSARDVKIMATYVCNLHRRSVRSVHEPDNTFRSPDASSLV
DVSNRPWRSSPCGNTGWDCTPEFDSHPIQSSVVDGDDTTSITLADIQRWLSANGYHRMFLPENSNNLWQAKIGI
INENADINQDTENGFGRVNYAIPFLGDLRTNMAKRDKELDWPVVAPRSLGDVRKNCCLRECSVEDFYGACS

***Scylla olivacea*** GDRN01043427.1
MRVVVGVVAVVALVVAVAEGDSLMTRSKKYKLCTARDINMFVTQLCSGALTRMRVGRHNSHSAPGILTIRIPPR
DVRGPVVSPAEIGSSYEFPSYRENLLDTQKYSPSELDAVDAPYLRPVSYWEGQSWNNYKESSAESELPSEYYGN
ALSPRNKKWWNDQWNAGEAGSAMIGLRRKRTTFGDIRQRCCTDGCYASELQSLCA

***Stenopus hispidus*** Trinity
MRAAGVVAVLVLVVVLVCVVVETRPPRIRRGLKICSARDVKFMATFVCNLHRRSVRSVVQEEEEEEEDDGDSDD
GDMGSPLDFKAIEWLAMMNPSQAYLQHWCGHSGVPCNPQRSFQRQQQAAPEDQQLDLRHRITPYVFNDWLSQIG
YSTNTSPNSWRRSEENDEMDDDALANTDTKSPLTVPAQNAFRLKVSKRDKEVDWPSMVPSTLGDIRKSCCVREC
SAEDFYGACV

### **Table S4.** Predicted decapod gonadulin 2 precursors.

Species name and identifier are indicated; Trinity means that the sequence obtained from Trinity generated transcripts from public SRAs and Artemis that the sequence is predicted from either a draft genome or genomic SRAs. In yellow the predicted signal peptide, in blue the propeptide that will produced after removal of the signal peptide. Possible proteolytic cleavage sites are highlighted in black and cysteine residues in red. See note about processing of the gonadulin precursor in the main text.

***Carcinus maenas*** GFYW01079908.1 partial
MKKVAEVVMMVMMIVTVGAQEDKLVTTCSQRDLRKAINNICSRRRQRSIRSSPSISSFSFSHSSFPQSFLSFPH
SSSSSLSFPYSYLXXXXXQPRTHTSHQQDIPEGHRKTYLLILASLKGRETMDVAWRGVEGMEGRVGQAKNVQGR
RMDAIGRTELKSGWRMGKNKWWVNKNAIKRHFLTKRDVSTLRILRRRCCLQGCRESELISACR

***Cherax quadricarinatus*** Artemis
MGGFETMMMVIVMVMVVVESHPSRTKRSGRICSSREVLQAANVVCNLTRRSAPASVFSGTLSSASRGVTAVASG
GVTAGDAFGDAYSGVTLSTPSPRNLDWLYGRSYKKPFRVTSPSPETRFILNNGISNRKRASTVVNLGTTGVVNS
WKNLIGTRESDPTGVTGRIRNRQNPVVKRHLNAETLTLRELRIICCVRECTIQDFLGACS

***Eriocheir sinensis*** Trinity incomplete
MRGVDMAVVLKVMMVVMVVAADEKDSEERVTVCSQRDFQHTITNICRSRRRRTAWSSSSSSVPSLSSSSSILSS
FPSSSTSSLSFSSS

***Jasus edwardsii*** GGHM01012768.1
MRTVGALVLVVVLAAAMVETRPYEETKSYKICTSRDVKVMANYVCNLHRRRRSIFSLNDARDTYGVPELLFESR
SRRLALPQHWRPQDHTATQDAEPGMGWINVSRRDPSFLQFTRDRRQVLLGEIRKQCCVAGCTPRDFYGACQ

***Macrobrachium nipponense*** Trinity
MRTFLLVPLIPTVLVLPLVYAGGPTDDANWPRSRRICSPRQVDLRAYEACHLTKRRSVDGAEEGQEVAGDSHLG
LTSNQARIPNSPSSVTSTKIFPPTKTHRPTRSEDDFGQFKRRMKVLKRRSSPPFKRQVTDVPDWRELNFTTYEE
VRLFCCRRSCPPEAYYGIC

***Macrobrachium rosenbergii***
MRRFLLVPLMITILVPGLACGGGPIDDANWPRSRRICSPRQVDLRAYEACHLTKRRSVDSVEEGQQTAGGSHLG
LTSNQAQAPNSPSVIGTRTIPSNKAHRPKRSTDDFGQFKRRMKALKRQGSLPFKRQVTDVPDWRELNFTTYEEV
RLFCCKRSCPPEAYYGIC

***Macrobrachium tolmerum*** GHDQ01053345.1
MRTFLLVPLMILTFLVPALVSGGGPTGNANWPRARRICSPRQVDLRAYEACHLTKRRSVDSAEEGRETSGGSHL
GMTSNQGHDSNSPSVSNTKTIPATKAHRPKRFADDFGQFKRRMKALKRQSSPPFKRQVTGVPDWRELNFTTYEE
VRLFCCRRPCPPEAYYGIC

***Menippe nodifrons*** from SRR8787123
MKGITAVVVTMVVTTVAGEGNQLVKMCSQRDFRSTLSDLCSRRRQRSVWSFPSSTSFSQSSLSFPHFSSSMSSK
SYPYSSDISSFSIPFSSSSSALTHLGIKGRPRTHPRRLQDTFGGHTKTYLLLPASLKGHGAVGVAWRGAEGVVL
RRMGAGGSREDESGWKVGKGRGKWRAGNEVTKRHVLVKRTANTIADLRRRCCLHGCRENELISACR

***Ocypode quadrata*** from SRR8778597
MKGSNVAVGMMLMIAMMVAAEKENEQRVILCSQRDFQLTISNACSRRRRRSALTFPSSFLPPFSSSYYFSTILL
SSSSPFFSSSHSVSSFASFSYISPRSSTTSTSSSSFLPLFYHLDIKEPRNFPSERLANLQESPRGHTETLSLTP
LNQHGPENIAWGEREGIQALELRTGDLGVQMSERRTSIDEWRVKIIKKVSVDVWTVDIVKWMTGMGKVSLGMGS
WTRVSERVPKGDLARHLVKRRTRTVSELRQKCCLKGCRVSELLAVCK

***Palaemon carinicauda*** Artemis
MRILLQLSAMQTILVLVLIAAEGPSDNATWARSRRICSPRQVDIRAYEACHLTRRRSVANNEKRRPIGSHLGSL
SSSDNSVHSRKGKYANHILSVRPNQHGSFPEDFGQFKRRMKTLKKHSTSTSSNAVFKELEELAPKRYFKRQVTD
VPDWRELNVTTYEQVRLFCCRRPCPPEAYYGIC

***Paralithodes camtschaticus*** GHJC01008804.1
PLQLLLLVLELLLTKEGRAWPALRGRRSTSDLCTPREIRRIANDVCNIARRSIRTQLTPSHHETDWPLLRRTPN
GIPAGHEGTNNYRFSLPSPLTLSSDVRPGVHSLIRSNTKLSQVLVPPLSNTIKARLASQAMVWRLMLAGMVRRK
RLNRRQTQHQSTTMTLEELRETCCNAQCSEQDFLAACS

***Portunus sanguinolentus*** Trinity
MNVAATVVAAVVVVVVVVTAVEGDLRVTVCSQRDLRNTISNLCSIRRQRSIWSFLSSSSSSSSSSSSSHQTISF
PHTFTYLDSSTSSPSPSSSSSSLPTHLGIKGRPRTHNTQEDTTEGHTRTYPLILPSLVERLSVGLSLHGAEGVG
RHRVGAGGRRSRWRVGKNKWWVSKDKIKRHFLTKRRVRTLGDLRRRCCVHGCLESDLLSACI

***Portunus trituberculatus*** Trinity
MNVVAVVAVVVVMVVTVVEGDPRVTVCSQRDLRNTISNLCSIRRQRSIWSFLFSSSSPPSSSSSSHQSISFPHT
FTYSDSSTSSPSSSYSSSSSSSSSSSSLLTHLGIKGKFPRNQGRPRTHSTQEDTTEGLTRTYPLILPSLVGRGD
VGLSMHGAEGVGRHRVGAGGRRSRWRVGKNKWWVNKDAIKRHFLTKRKVRTLGDLRRRCCVHGCLESELLSACI

***Sagmariasus verreauxi*** AIU40994.1
MRTVGALVLVVVLAAAMVETRPYEETRSYKICTSRDVKVMANYVCNLHRRRRSVLSLDDARDNYGVPGLLLENR
SRRLALPQHWRPEDDTDTGNVSRRDPSFLQFTRIRRQVLLGEIRKQCCVHGCTPRDFYGACQ

**Table S5.** Predicted decapod gonadulin 3 precursors.

Species name and identifier are indicated; Trinity means that the sequence obtained from Trinity generated transcripts from public SRAs and Artemis that the sequence is predicted from either a draft genome or genomic SRAs. In yellow the predicted signal peptide, in blue the propeptide that will produced after removal of the signal peptide. Possible proteolytic cleavage sites are highlighted in black and cysteine residues in red. See note about processing of the gonadulin precursor in the main text.

***Macrobrachium nipponense*** Trinity incomplete

MKGIGYYLATAITALLFIRGTESAAHLFRVGRSIRVCTPRDVARIGHFMCKRSVLPAGEQQDGGTTIRQDERSL
EHREKSKDSNTFSSAESIERDAFWEESPDDAFHDNIYPFLELKGVXXXXVARKLQKRSNNRPRKFETIADIRKY
CCHDDCPIELFLCI

***Macrobrachium rosenbergii*** Trinity

MKGIGYYLAIAITALLFIRGTESAAQLFRVRRSIRVCTPRDVARIGHFMCKRSVLPTEGQDGAGNVRPDERSLE
HQERSKDSNTFSRAESSGRDAFWEESPEHAFHDIYPFLEFKRVTYYEGSGQIDGRYFDGSGHTEGSYFGGSGQI
EGSYSDGSGLIDGSYFDGSGQIDGSYFDGSGQIDGSYFDGRGHIDDSLTERKLQKRSNKWPRKFETIADIRKYC
CHHDCPIELFLAECA

***Palaemon carinicauda*** Artemis

MKSSGYLLATAVTALLLIRGTESGTQVFRNRRSIRVCTPRDVARIGHFLCKRSISSMDANKRGLDESVHNGDAT
EKEVGANSNLFSSMGAFEVADTNESKQDMLPEIFMRFSEPEGRGSPASDGDLENRKLQKRSSSWPRRKFETIGD
IRRYCCHHDCPIELFIAECA

***Palaemon serratus*** Trinity

MKAIGYLLATTVTALLFIKGTESATQVFRNRRSIRVCTPRDVARIGHFLCKRSISSLDANKGGPDEPVQKGDTT
EKEVGANSNLFSNTEPFGAADRNESKGDKLPEMFMRFSEPEGRRSPANGGDLEEKKLQKRSSPWPRRKFETIGD
IRKYCCHHDCPLELFIAECA

**Table S6.** Predicted decapod insulin precursors.

Species name and identifier are indicated; Trinity means that the sequence obtained from Trinity generated transcripts from public SRAs. In yellow the predicted signal peptide, in blue the likely mature peptide that will produced after cleavage by convertase at the proteolytic cleavage sites that are highlighted in black. Cysteine residues have been highlighted in red.

***Cancer borealis*** GEFB01034529.1

MKMKILLLVVVTAMQAGRTLGSPRTLPEGEAVSEGERRLCGWKLANELNRVCKGVYNKPTVSTNALFYLKERGG
KRVDLWPLGIELQFPPWTQAPADERRANQIIESHQPDKPMEQSLGESQRLVFLTGPEASQVVSGLPRVKRGLSA
ECCRKACSVSELAGYCY

***Carcinus maenas*** GFXF01207286.1

MKVQRQVLLLVMVTAMHAGITRSSPRTLPGQGLVTDGERRLCGWRLANELNRVCKGVYNKPTVSNNALFYLRGR
GVKRVDLMPLGMELEFSPWTQASADDLRSDQISESHTSHIPKQLPFLAEAEASRVVGGLPRVKRGLSAECCRKA
CSVSELAGYCY

***Charybdis feriata*** GGFD01070073.1

MKVVIFLLLVATAMQAGKTSGSPRTLSEGGLVREGERRLCGWRLANELNRVCKGVYNMPTVSSNALVYLKGRGG
KRVDLWPIGRELRYPSWTQAPSDDLLSGQHSEPRLLHKPMERSSSERQRLPVLTGAEASQVVGGSPRVKRGLSA
ECCRKACSVSELAGYCY

***Cherax quadricarinatus*** Trinity
MKVIVLVVGVVVWLVLGSTGSHPSNYPNSELELEPRRRLCGWRLANKLNSVCKGVYNKPQYADNLLYYRGRRVG
GKTGLQQQPEALKELFSDDIVMASRGHSTSTPPVLPIVSTDTAEQDNRELEDLLNMQLSQDPSQSASRENEDS
KTHLPFLTKMEALQMVRNRPRPKRGLSAECCQKVCTVSELVGYCY

***Eriocheir sinensis*** Trinity
MKVMVLLLVAAAAAAQPSKSRGPLKTLPGAGAVREAERRLCGWKLANELNRVCKGVYNKPTVSTNALFYRRERG
EGSVDFEDPVDVWPLMMELDFSPWTPAPPDVVRRGVLPGPPVVPASEDQRLPFLSGPQASQVVGRSRRVKRGLS
AECCRKACTVSELAGYCY

***Homarus americanus*** GEBG01059205.1
MRAFVVVIAVVVVVVLELGSSRASRRTYPTSEEEPRRRLCGWRLANKLNLVCKGVYNNPGSTGNYLFYRSRRDG
ESEPGLPPEKYLDLLADPEEERGLRHHYLTSSQQASEDTPSEENEAPGSFFGSLSPQDLPHQSAVQEDEASSVH
FPFLTEEEASQMVRVRPRSKRGLSAECCRKVCTVSELVGYCY

***Jasus edwardsii*** GGHM01050027.1
MKAIVVLVAAAAAVVQLGSGRDSVQSSFAMNEAEEEPQRRLCGWRLANKLNKVCKGVYNKPGSTTNLYYYYRDR
RAGGGSEPVPPPDESLAAPVAARVSKYRPLPPRESLPSYTSDVVNSWGDESTGAVLQPDLPQELPLQLAAHRAG
DLQGLRFPFLTKTEASQVLHTQQRSKRGLSAECCRKVCTVSELVGYCY

***Macrobrachium australiense*** GHDT01024471.1
---SSSSLAQVGSSDLGEEEKPLRRLCGWRLANKLNQVCKGIYNKPTVTNNDLFYRSMRGGGPLYDFR
PKTPEISRDDDIRYHFPLTDDYYYFYSSDARESDGLPLEGYVQEGKESPPFLSRQEAAQMFKAHPRSKRGLSAE
CCRKACRVSELMGYCQ

***Macrobrachium nipponense*** GCVG01033096.1
MRTLNAVLLLTLILQGGVGSPRGSSPSSSSLAQVSGSSSDLGEEEGKPLRRLCGWRLANKLNQVCKGIYNKPTV
TNNDLFYRSMRGGGPLYYEFRPKTPEVSDDFRYHFPKTDDSYYYYYYSGGRETDDGGLPPGGYALEGQERSPFL
SKQEASQMFKAHPRSKRGLSAECCRKACRVSELMGYCQ

***Macrobrachium tolmerum*** GHDQ01067937.1
MRTLNAVLLLTLILQGGVGSPRGSPSSSSLAQVGSSDLGEEEKPLRRLCGWRLANKLNQVCKGIYNKPTLTNND
LFYRSMRAGGPLYDSRPKTPEISRDDDFRYHFPMTDDDYYYYYFYSSDGRESDGSPSEGYVREGKESPPFLSRQ
EAAQMFKAHPRSKRGLSAECCRKACRVSELMGYCQ

***Menippe nodifrons*** Artemis
ILLLVVVTAMQAGKTRGSPRTLPGGGAAREGVRRLCGWRLANELTRVCKGVYNKPTVSTNALFYLKERGGKRVD
LWPLGMKLELPSRTQAPADELRIDQISEPQPAYPPVERPPSESRQLPFLTGAEASRVVGGSPRVKRGLSAECCR
KACSVAELAGYCY***Neocaridina denticulata*** GGXN01032148.1
MKALIVLLLLNQLLGGAYASPHDSSSSSSLSLTSSLSEMTDLDEEERPLRRLCGWRLANKLNEVCKGIYNKPGP
TSNDLFYRTLRGSPYSDISSAYEGVSGPLMAKTTQTDYDALLGARKSPSIGDLMSDMKETAEPGQKLELFDQVL
PYVEAAAFHIDQRAPSMSSVVEELRKSVISSGETSFPFWTRKEASQMLKLHPRSKRGLSAECCRKACRISELMG
YCQ

***Palaemon varians*** GFPG01026539.1
MRTFNTVFLLTLILHCGAVSPRGSSSVASLSLASSLSQSSDLEAKEKPMRRLCGWRLANKLNQVCRGIYNKPSG
TSNDLFYRSMRGESLNDFKPDIGIETDLSSTKDTKLDLIDSPKNEKHIPLPGRRVPPETSKDNFRYLFPTAEDY
NYYLEGRGIDGISPEGYVRKEKERFPFLSSEVASQMFLSHPRAKRGLSAECCRKACKISELMGYCQ

***Palaemon***
MRTLNTVFLLTLILQGGAVSPRGSSPVSSLSLASSLAQVSDLEVEEKPMRRLCGWRLANKLNQVCRGIYNKPSV
TSNDLFYRSMRGEPLNDFKPEFGIETDVSSTKDTQVGLIDSLKNEKHIPSSGRRVPPEILNGNFRYLLPTTEDY
KYYFEGQDNDGISPEGYVRKEKERFPFLSSEVASQMFLSHPRSKRGLSAECCRKACRISELMGYCQ

***Panulirus argus*** genome reads
MKVVVALIAAAAVVVKLGSGRDSQSSFPVSEVEEEPQRRLCGWRLANKLNKVCKGVYNKPGSTRNLYYYYRDRR
AGGGSEPVPPPDESLAAPSAARVFRYRPLSPRHLLRPYTSDVIDSWAEGDRSSGAIFEPRLPQELPLHSAQYNE
DGGIQSGQSPFLTEAEASQVVRTRQPSKRGLSAECCRKVCTVSELVGYCY

***Penaeus vannamei*** XP 027231326.1
MKIAIAVFLALVCLQSGCSWMTDLDTSREPQRRLCGWKLANKLNSVCKGVYNKPGPMSNSLYYRHRRAKTRPRI
GDFSASLPRTTDSRQSPRGDAGLWDLSYPPNPPPSPPTPPTTSPSPLTAPTPSSSPPQPRAPPPSATDAFDQLL
LHNALLSSGVPFPFLTEAEASQMLKEAPRRKRGLSAECCRKACSVSELVDYCY

***Portunus sanguinolentus*** GFZC01032047.1
MKVLLLLVVTAMQTGRVRGSPRTLPEGGLVKQGERRLCGWRLANELNRVCKGVYNVPTVSTNALFYLKGRGGKR
VDLWPVGGREQHFPSRTQAPAEDLRAGQLSETHLFHRPADQPTGERQRLPLLTGAEASQVLVGRSPRVKRGLSA
ECCRKACSVSELAGYCY

***Portunus trituberculatus*** GFFJ01005788.1
MKVVVLLLVVVTAMQTGRVRGSPRTLPEGGLVKQGERRLCGWRLANELNRVCKGVYNVPTVSTNALFYLKGRGG
KRVDLWPVGGREQQFPSRTHAPADDLRASHLSEPHLFQRPADRPTGERQRLPLLTGAEASQVVVGRSPRVKRGL
SAECCRKACSVSELAGYCY

***Procambarus clarkii*** GBEV01062121.1
MQAPVVVVVVVVALLDLGSSGASQDTYTTSHPEGEPGRRLCGWRLANKLNRVCKGVYNNPRSTNNYLYYRGRRV
DERRTPEQPANELLDVLPDELVDMRGPRLRLPQPTVPGPPYVSRGPAYDSRDPAYVSRGPPYVSRGPHPPPPPG
EQDSGAQRAFLTLKEAAQMLKTQPRHKRGLSAECCQKVCTVSELVGYCY

***Scylla olivacea*** GDRN01009614.1
---MQTGRTRGSPRTLPVGGLVREGERRLCGWRLANELNRVCKGVYNMPTVSSNALFYLKARGGKRVD
LWPVGRELQFTSWTQAPVDDLRAGQLSEPRLLHRPVSSSEYQRLPLLTGAEASQVVGGSPRVKRGLSAECCRKA
CSVSELAGYCY

***Stenopus hispidus*** Trinity
MVQLVALLGLLLQLLVGLSCASATMTQDDLSQSTGKGNQGLLRNTRDRQFMHTHSRERHEGSDVGVVDRKIQRL
CGSKLASKLNRVCKGVYNKPEPRYNHLYYRGRRGKSFSDVVRPPQQLTYGKSTSESESEVDNVGSQYTHGEEGR
EILPVSDVTQWEDSRDPPADMDLIPINGRSHLTPDDFYSNDDFEIFPPNDDHLRLAETDKDKFKFKNSIDQMLL
STYGKILSTSGQFPFLTEKDALKVLKNRPRRKRSLIAECCRKPCSISELVGYCD

**Table S7.** A possible correlation between the expression of relaxin and vitellogenin in the ovary of *Penaeus monodon*.

| SRA | Spots | Vitellogenin | | Relaxin | | mRNA source |
|---|---|---|---|---|---|---|
| | | | | | | **Fed pellets** |
| SRR9031901 | 54400454 | 16 | **.29** | 70 | **1.3** | before eyestalk ablation at week 0 |
| SRR9031902 | 61068478 | 30 | **.49** | 86 | **1.4** | before eyestalk ablation at week 4 |
| SRR9031899 | 52693150 | 38 | **.72** | 42 | **.8** | after eyestalk ablation at day 1 |
| SRR9031900 | 48043587 | 50 | **1.04** | 422 | **8.8** | after eyestalk ablation at day 4 |
| SRR9031905 | 59848970 | 1288 | **21.52** | 71 | **1.2** | after eyestalk ablation at day 9 |
| | | | | | | **Fed polychaetes** |
| SRR9031906 | 59926910 | 15 | **.25** | 272 | **4.5** | before eyestalk ablation at week 0 |
| SRR9031903 | 48830634 | 321 | **6.57** | 1230 | **25.2** | before eyestalk ablation at week 4 |
| SRR9031904 | 57947220 | 2008 | **34.65** | 3067 | **52.9** | after eyestalk ablation at day 1 |
| SRR9031907 | 117972925 | 24804 | **210.25** | 1723 | **14.6** | after eyestalk ablation at day 4 |
| SRR9031908 | 61527503 | 19429 | **315.78** | 712 | **11.6** | after eyestalk ablation at day 9 |

Spearman value: 0.539;  $P < 0.10$ one sided

Data from :

Sittikankaew, K., Pootakham, W., Sonthirod, C., Sangsrakru, D., Yoocha, T., Khudet, J., Nookeaw, I., Uawisetwathana, U., Rungrassamee, W., Karoonuthaisiri, N., 2020. Transcriptome analyses reveal the synergistic effects of feeding and eyestalk ablation on ovarian maturation in black tiger shrimp. Sci Rep. 10:3239. doi: 10.1038/s41598-020-60192-2.
